# Müller glia–mediated regeneration restores neuronal diversity and circuit organization in the adult zebrafish retina

**DOI:** 10.64898/2026.03.15.711785

**Authors:** Mikiko Nagashima, Sherine Awad, Lu Jiang, Lara S. Rappaport, Zachary Flickinger, Peter F. Hitchcock, Thanh Hoang

**Author notes:** These authors contributed equally to this work (Mikiko Nagashima, Sherine Awad).

## Abstract

The ability to regenerate neurons with appropriate identities and circuit connectivity is a fundamental challenge in regenerative biology. Unlike mammals, adult zebrafish robustly regenerate retinal neurons after injury through the reprogramming of Müller glia. However, the extent to which regenerated neurons faithfully reconstruct molecular identity, cellular diversity, and circuit organization remains unclear. Here, we combined inducible lineage tracing, single-cell RNA sequencing and high-resolution morphological analysis to define the molecular identities and structural organization of regenerated neurons following photoreceptor-selective light lesion or NMDA-induced inner retinal injury. Both injury paradigms regenerated all major retinal cell classes, although the relative abundance of regenerated cell types reflected the pattern of neuronal loss. Across major neuronal classes and subtypes, regenerated neurons largely reestablished endogenous molecular identities, with the residual transcriptional differences primarily reflecting ongoing maturation. Regenerated amacrine and bipolar neurons recovered subtype diversity, characteristic dendritic morphologies, and laminar organization. Regenerated retinal ganglion cells likewise restored broad molecular diversity and appropriate retinotectal projections, while a subset underwent microglia-mediated refinement. Together, these findings demonstrate that Müller glia-mediated regeneration largely reconstructs neuronal identity, cellular diversity and key features of retinal circuit organization, providing insights into understanding how complex neuronal tissues are rebuilt after injury.

## Introduction

The development of the nervous system, such as the vertebrate retina, requires the orchestrated differentiation of multipotent progenitors into diverse types of neurons and glia, and their organization into local and long-range circuits(Agathocleous and Harris 2009; Cepko 2014; Götz and Huttner 2005). These processes are governed by temporal and spatial patterning that regulate progenitors’ transition from proliferation to cell fate specification and circuit assembly(Götz and Huttner 2005; Livesey and Cepko 2001). This hierarchical organization, from molecular identity to circuit architecture, is essential for visual processing. Regenerating neurons in the adult neuronal tissues after injury presents significant challenges, as neurogenesis must be reactivated within mature tissue where architecture and connectivity are already established, and newly generated neurons need to adopt appropriate subtype identities and integrate into existing circuits.

Although the adult mammalian central nervous system possesses a limited spontaneous regenerative capacity(Ramón y Cajal 1991; Richardson et al. 1980), recent advancements have demonstrated that endogenous glial cells possess latent neurogenic potential and can be induced to generate new neurons. For instance, forced expression of proneuronal transcriptional factors can reprogram subsets of astrocytes into neurons in the adult brain (Götz et al. 2015; Bocchi et al. 2022). Likewise, mammalian retinal Müller glia, which typically remain quiescent and undergo reactive gliosis following injury, can be induced to generate new neurons(Jorstad et al. 2017; Todd et al. 2022; Yang et al. 2025; Hoang et al. 2020; Le et al. 2024). These studies have established the feasibility of glia-derived neuronal regeneration. However, two major barriers remain: the limited diversity of regenerated neurons and the uncertain extent to which those neurons acquire the mature molecular, morphological and circuit-level properties required for functional restoration.

Unlike mammals, adult zebrafish retina possesses a remarkable capacity to regenerate retinal neurons through reprogramming of Müller glia(Lenkowski and Raymond 2014; Karl and Reh 2010; Hamon et al. 2016);(Bernardos et al. 2007). Following injury, Müller glia de-differentiate, re-enter the cell cycle, and divide asymmetrically to generate multipotent Müller glia-derived progenitors that can produce all major retinal cell types (Nagashima et al. 2013; Lenkowski and Raymond 2014); (Powell et al. 2016; Lyu et al. 2023; Lahne, Brecker, et al. 2020). While considerable progress has been made toward understanding the molecular mechanisms that initiate Müller glial reprogramming (Lahne, Nagashima, et al. 2020; Hoang et al. 2020; Nagashima and Hitchcock 2021; Jui and Goldman 2024), these studies have largely focused on the early signals that activate Müller glia and the transcriptional programs of the resulting progenitors. Considerably less is known about the fidelity with which regenerated neurons reconstruct retinal organization. Specifically, it remains unclear whether regenerated neurons faithfully recover endogenous molecular identities, regenerate the broad spectrum of neuronal subtypes, restore characteristic cellular morphologies and laminar organization, and rebuild appropriate long-range neural circuits.

Here, we address these open questions by combining inducible genetic lineage tracing, single-cell RNA sequencing and high-resolution morphological analysis to comprehensively evaluate Müller glia-derived regenerated neurons in adult zebrafish retina. By comparing photoreceptor-selective light lesion and NMDA-induced inner retinal injury, we found that both injuries induced Müller glial reprogramming to generate all major retinal cell classes, while injury context influences the relative abundance of regenerated neurons. Transcriptomic analysis showed that regenerated neurons largely re-establish endogenous cell type and subtype diversity. Morphological analysis further demonstrated that regenerated neurons restore characteristic cellular morphologies, lamina-specific dendritic stratification, and long-range axonal projections. Together, these results demonstrate that adult zebrafish retinal regeneration restores complex neuronal identities at molecular and structural levels, providing insight into how regenerative neurogenesis achieves circuit reconstruction in the mature nervous system.

## Results

### 1. Injury-inducible genetic lineage tracing identifies Müller glia-derived neurons

Before assessing the identity of regenerated retinal neurons, we first validated the cell type specificity of our injury paradigms. Since previous work shows that prolonged light exposure damages inner retinal neurons in addition to photoreceptors(Lyu et al. 2023), we performed TUNEL apoptotic assays to determine if our brief, high-intensity light lesions also damage inner retinal neurons. TUNEL assays performed immediately following light exposure revealed a robust accumulation of apoptotic cells within the outer nuclear layers (ONL) that peaked between 1 and 2 days post lesion (Fig. S1A,B). In contrast, no TUNEL+ cells were observed in the inner nuclear layer (INL) or ganglion cell layer (GCL) (Fig. S1A,B), and the number of HuC/D+ neurons remained unchanged (Fig. S1C,D). Conversely, intravitreal NMDA injection selectively ablated inner retinal neurons without detectable damage to photoreceptors in the ONL (Fig. S1E,F). These data confirm that our two injury models selectively ablate distinct retinal neuronal types.

To unambiguously distinguish regenerated neurons from endogenous cells, we employed inducible Cre-loxP–based genetic lineage tracing. We generated a transgenic zebrafish line by crossing *Tg(mmp9:CreERt2)* with T*g(Olactb:loxp-dsred-loxp-egfp)* reporter (Fig. 1A)(Ando et al. 2017). In this system, the Medaka actin promoter drives ubiquitous dsRed expression in all retinal cells under normal conditions. Following a retinal injury, *mmp9* promoter activity is specifically induced in reprogrammed Müller glia and their progenitors(Silva et al. 2020; Lyu et al. 2023). Administering 4-hydroxytamoxifen (4-OHT) after a retinal lesion induces CreERt2-mediated recombination selectively in activated Müller glia, permanently switching the reporter expression from dsRed to eGFP in lineage-traced cells (Fig. 1A). As expected, unlesioned retinas showed strong dsRed signal without detectable eGFP (Fig. 1C,D).

**Figure 1.**
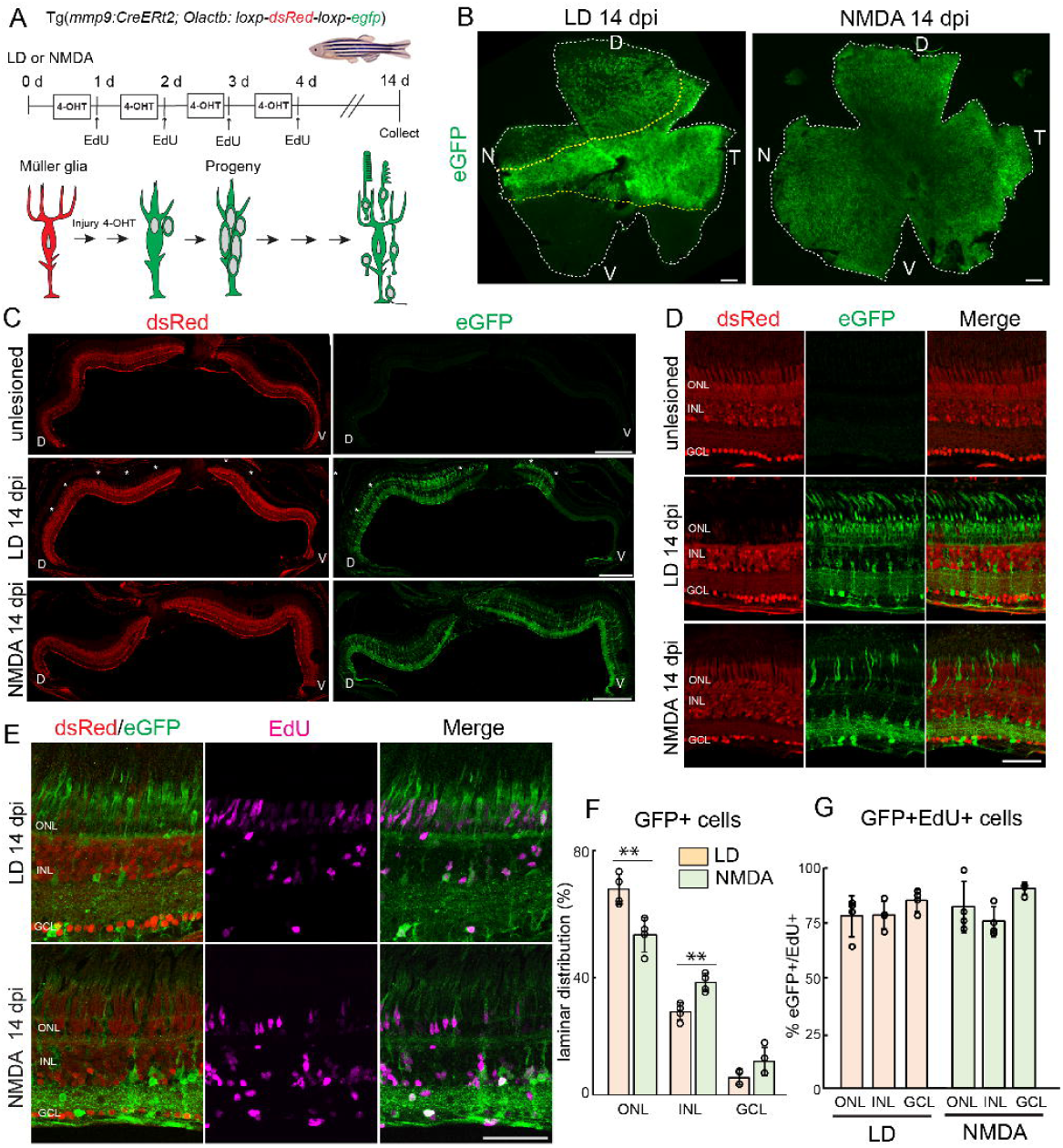
Injury-inducible lineage tracing specifically labels Müller glia-derived progeny. (A) Experimental paradigm of retinal lesions, 4-hydroxitamoxifen treatment, and EdU injection to label Müller glia-derived progeny using inducible transgenic lineage tracing line, *Tg(mmp9:creERT2; Olactb:loxp-dsRed-loxp-eGFP)*. (B) Flat-mount retinas immunostained for eGFP at 14 days post light lesion and NMDA injury. Yellow dotted lines indicate regions of photoreceptor damage. (C,D) Retinal cross sections immunostained for eGFP in unlesioned, 14 days post light-lesioned and NMDA-injected retinas. Asterisks indicate regions of photoreceptor damage. (E) Retinal cross sections immunostained for eGFP and EdU at 14 days post light and NMDA damage. (F) Laminar distribution of eGFP+ cells in the outer nuclear layer, inner nuclear layer, and ganglion cell layers at 14 days following light lesion and NMDA injury. (t-test, ONL: p=0.0078; INL: p=0.0018; GCL: p=0.0878)(G) Quantification of the proportion of eGFP/EdU double positive cells from total EdU+ cells in light-lesioned and NMDA-injected retina. Scale bars: B,C: 200um; D,E 50 um; 4-OHT: 4-hydroxitamoxifen; EdU: 5-Ethynyl-2-deoxyuridine; LD: light damage; NMDA: N-methyl-D-aspartate; dpi: days post injury; D: dorsal; V: ventral; N: nasal; T: temporal; ONL: outer nuclear layer; INL: inner nuclear layer; GCL ganglion cell layer.

To establish the specificity and inducibility of the *mmp9:CreERT2* system, we performed two control experiments. Treatment with 4-OHT in the absence of injury did not induce eGFP expression (Fig. S2A), confirming that the *mmp9* promoter is inactive in uninjured retinas, and that CreERT2 does not exhibit leaky activation. Conversely, light lesions without 4-OHT administration did not induce eGFP despite robust endogenous mmp9 upregulation (Fig. S2B), demonstrating that CreERT2-mediated recombination strictly requires activation by the 4-OHT ligand. Together, these control experiments demonstrate that the *Tg(mmp9:CreERt2; Olactb; loxp-dsred-loxp-egfp)* system shows no detectable background recombination and provides tightly regulated, injury-dependent labeling of regenerated retinal neurons.

We next characterized the temporal dynamics of activated Müller glial and their lineages following a light lesion. This paradigm was selected due to its well-defined and highly synchronized response of Müller glia(Vihtelic and Hyde 2000). At 3 days post injury (dpi), eGFP+ cells appeared as radial columns spanning the entire thickness of the retina (Fig. S2C). These eGFP+ cells co-expressed the Müller glial marker, Glutamine synthetase (GS), confirming selective labeling of injury-activated Müller glia (Fig. S2C). At 3 dpi, Müller glia–derived progenitors appear as elongated nuclei associated with radial processes. The intensity of the eGFP signal in Müller glia-derived progenitors was relatively weak, likely due to delayed eGFP reporter accumulation. Between 3 to 5 dpi, Müller glia-derived progenitors proliferate and migrate into different retinal layers(Bernardos et al. 2007; Nagashima et al. 2013). By 7 dpi, subsets of eGFP+ cells in the outer nuclear layer (ONL) expressed cone (Zpr1) and rod (Zpr3) photoreceptor markers (Fig. S2D,E), indicating progression toward photoreceptor differentiation. These regenerated photoreceptors displayed immature morphologies, including shortened outer segments, consistent with ongoing maturation. These data demonstrate that the mmp9-driven lineage tracing system faithfully captures the progression from Müller glial activation through progenitor proliferation to neuronal differentiation.

We next examined regenerated neurons at 14 dpi, and compared regenerative outcomes between light lesion and NMDA damage. Analyses of retinal flat-mounts revealed that eGFP+ regenerated neurons spatially corresponded to regions of cell death, with light lesions inducing photoreceptor death within a horizontal band spanning the nasal-temporal axis, with partial loss in the dorsal retina and sparing of the ventral retina (Fig. 1B,C; (Qin et al. 2009; Nagashima et al. 2013). The distribution of eGFP+ regenerated neurons precisely mirrored these patterns, with strong eGFP label in the central band, sparse eGFP dorsally, and no detectable eGFP in the ventral retina (Fig.1B,C). In contrast, NMDA injury resulted in a pan-retinal distribution of eGFP+ cells (Fig. 1B,C), consistent with widespread loss of the inner neurons. Consistent with the targeted damage of specific cell types (Fig. S1), light-lesioned retinas showed a preponderance of eGFP+ cells in the ONL along with additional labeled neurons in the inner nuclear layer (INL) and ganglion cell layer (GCL) (Fig. 1C,D). In contrast, NMDA-injected retinas exhibited prominent eGFP labeled cells within the INL and GCL and sparse labelings in the ONL (Fig. 1C,D). These observations confirm that the *Tg(mmp9:CreERt2; Olactb; loxp-dsred-loxp-egfp)* lineage tracing system faithfully marks regenerated neurons in spatial patterns that reflect the known patterns of cell death.

Finally, to verify that eGFP+ neurons originate from proliferating Müller glia, we performed EdU incorporation assays (1–4 days post injury; Fig. 1A). At 14 days post injury, 75.6% (light lesion) and 77.7% (NMDA) of eGFP+ cells were EdU+, with comparable proportions across retinal layers (Fig. 1E-G). EdU+/eGFP-cells likely represent activated microglia or rod photoreceptors derived from eGFP-rod precursors(Stenkamp 2011; Nagashima et al. 2013; Mitchell et al. 2018). The high fraction of EdU+/eGFP+ cells confirms that lineage-traced neurons arise from proliferative Müller glia–derived progenitors.

### 2. Müller glia regenerate all major retinal cell types with injury-dependent differences in neuronal output

To determine how precisely neuronal cell types and their molecular identities are restored following damages to specific retinal cell populations, we performed single-cell RNA sequencing (scRNA-seq) on FACS-isolated eGFP+ cells at 14 days after either a light lesion or NMDA injury (Fig.2A). After cell filtering and quality control, we retained 11,214 cells from light-lesioned retinas and 15,424 cells from NMDA-injected retinas (Fig.2B). For unlesioned controls, we incorporated our previous scRNA-Seq datasets comprising 134,237 cells (Lyu et al. 2025). Unsupervised clustering and annotation using established marker genes identified all major retinal cell classes across all samples, including both injury paradigms (Fig. 2C,D; Table1). Although light lesions selectively ablate photoreceptors whereas NMDA preferentially eliminates inner retinal neurons (Fig. S1), for each injury paradigm all major retinal cell types were generated (Fig. 2B,C). This observation was validated using immunohistochemical analysis of light-lesioned (Fig. 2F,G) and NMDA-injected retinas (Fig. S3). In both injury paradigms, eGFP+ rod (Zpr3+) and cone (Zpr1+) photoreceptors, bipolar cells (Cabp5+), amacrine cells (HuC/D+), horizontal cells (identified by position and morphology), and RGCs (Rbpms+) were detected (Fig. 2F,G; Fig. S3). These results demonstrate that neuronal output from Muller glia-derived progenitors is not restricted to the ablated cell types. Instead, Muller glia-derived progenitors retain broad neurogenic competence and generate all major retinal cell classes regardless of injury context.

**Figure 2.**
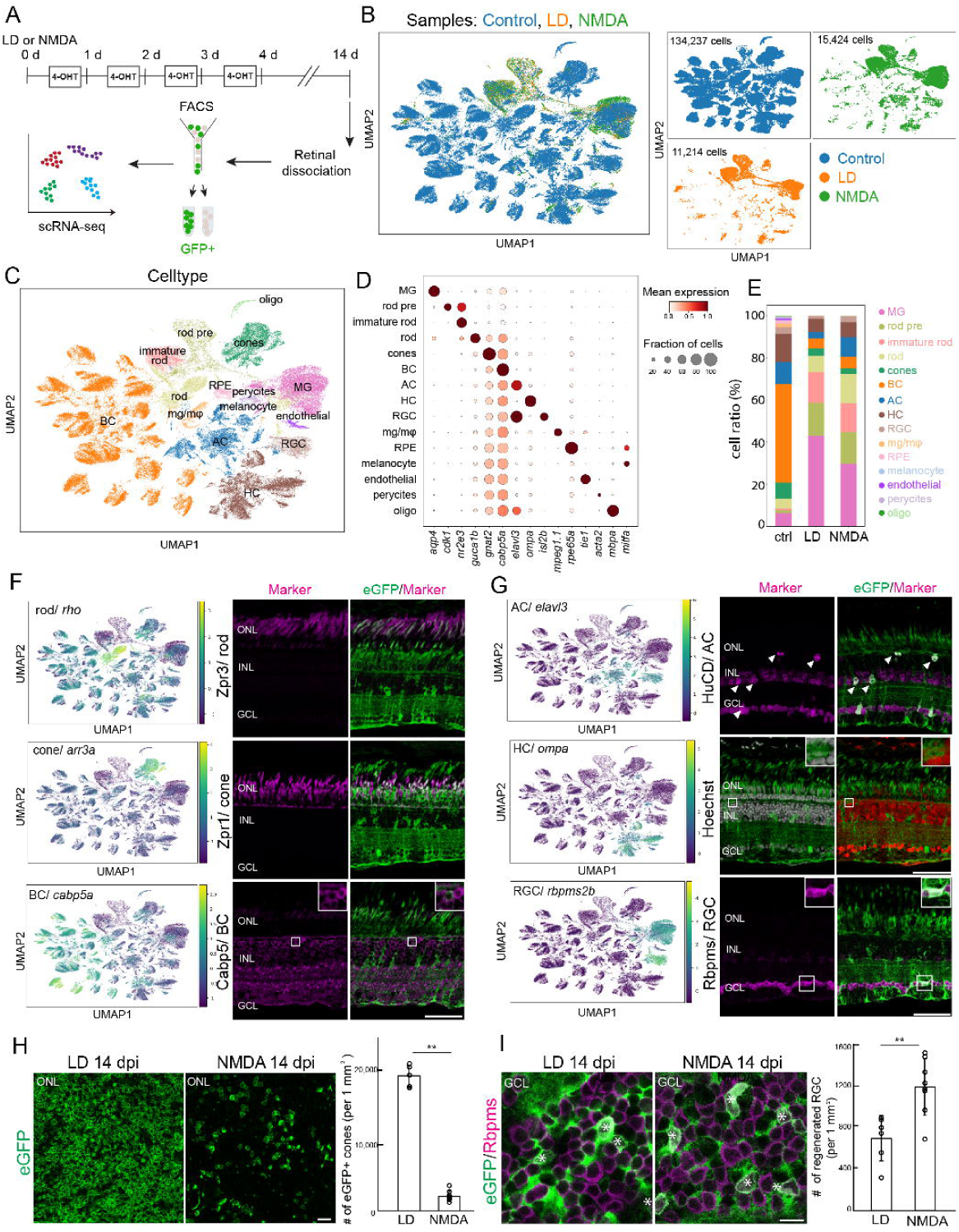
Regeneration restores all major retinal classes across injury paradigms. (A). Experimental design and scRNA-seq profiling of FACS-isolated eGFP+ cells after injuries. (B) UMAP plots showing the single cell distribution of control, light-damaged, and NMDA-injected samples. (C) Cell type annotation on UMAP plot across all samples. (D) Dot plot of selected markers of major retinal cell types. (E) Quantification of the ratios of retinal cells in control, light-damaged, and NMDA-injected samples. (F) scRNA-seq mRNA expression and representative immunohistochemistry for rod photoreceptors (*rho*/ Zpr3), cone photoreceptor (*arr3a*/ Zpr1) and bipolar cells (*cabp5a*/ Calbp5a) at 14 days post light-damaged retinas. (G) scRNA-seq mRNA expression and representative immunohistochemistry for amacrine cells (*elavl3*/ HuC/D), and RGCs (*rbpms2b*/ Rbpms), and Hoechst-labeled horizontal cells (*ompa*) at 14 days post light lesion. (H) Representative immunostained images of flat-mount retina and quantification of eGFP-labeled cone photoreceptors in the outer retinal layers at 14 days post light or NMDA-injection (p=0.0001). (I) Representative immunostained images of flat-mount retina and quantification of eGFP and Rbpms at 14 days post light or NMDA damage (p=0.0035). Asterisks indicate eGFP+/Rbpms+ regenerated RGCs. LD: light lesion; NMDA: N-methyl-D-aspartate; dpi: days post injury; BC: bipolar cells; AC: amacrine cells; HC: horizontal cells; ONL: outer nuclear layer; INL: inner nuclear layer: GCL: ganglion cell layer. Scale bars F,G: 50 um; H,I 10um.

We next compared the relative abundance of regenerated neuronal types between the two injury models and found a distinct difference in the proportion of cell types generated (Fig. 2E). Relative to a light lesion, NMDA injury resulted in a preponderance of inner retinal neurons, including amacrine cells (LD 3.05% vs NMDA 9.23%) and RGCs (LD 1.4% vs NMDA 2.96%) (Fig. 2E). The relative abundance of cone photoreceptors is comparable between the two injury paradigms (LD 3.32%; NMDA 2.75%), and NMDA samples contained a high frequency of rod photoreceptors (LD 7.9%; NMDA 13.74%) (Fig. 2E). To more accurately quantify the regeneration of photoreceptors, we analyzed regenerated cones in flat-mount retinal preparations. This analysis prioritized cones because they arise directly from Müller glia-derived progenitors(Lenkowski and Raymond 2014), and their well-characterized profiles allow for highly accurate cell counting(Nagashima et al. 2017, 2020). As expected, light-lesioned retinas exhibited a significantly greater number of eGFP+ cone photoreceptors compared to NMDA-injected retinas (Fig. 2H). Likewise, quantification of eGFP+/ Rbpms+ RGCs in flat-mount retinas showed that NMDA injury produced approximately twice as many eGFP+/RGCs as light lesions (Fig. 2I). These data suggest injury context does not confine the fate of Müller glia-derived progenitors to the neuronal classes that were lost to injury, but it does influence the relative abundance of their regenerated progeny.

### 3. Regenerated neurons recover cell type-specific transcriptional identities

To determine whether regenerated cells acquire the appropriate retinal cell identity, we performed transcriptomic comparison between regenerated cells and native retinal neurons. Müller glia showed the highest similarity to endogenous Müller glia in both light lesioned (LD: 0.87) and NMDA-injected (NMDA: 0.87) samples (Fig. 3A,B), suggesting that transcriptional changes associated with reprogramming are transient, and their glial identity is largely preserved after regeneration. Among regenerated neurons, each cell type exhibited the highest similarity to its corresponding endogenous counterpart with cones (LD: 0.85; NMDA: 0.86), RGCs (LD: 0.85; NMDA: 0.87), amacrine (LD: 0.83: NMDA: 0.85), and horizontal cells (LD: 0.83; NMDA: 0.82) (Fig. 3A,B), indicating that regenerated neurons largely re-established their native molecular identities. Regenerated rods exhibited slightly lower similarity (LD: 0.77; NMDA: 0.77) than others and displayed nearly equivalent similarity to immature rods (LD: 0.75; NMDA: 0.77), consistent with ongoing maturation at 14 dpi. Regenerated bipolar cells showed comparatively lower similarity scores (LD: 0.65; NMDA: 0.66), which may reflect their relatively limited subtype diversity (see next section) and/or ongoing maturation. Notably, corresponding regenerated cell types from light-lesioned and NMDA-injected retina displayed the highest similarities (>0.94) (Fig. 3C), suggesting that the transcriptional signatures of Muller glia-derived neurons are pre-programmed and independent of injury context.

**Figure 3.**
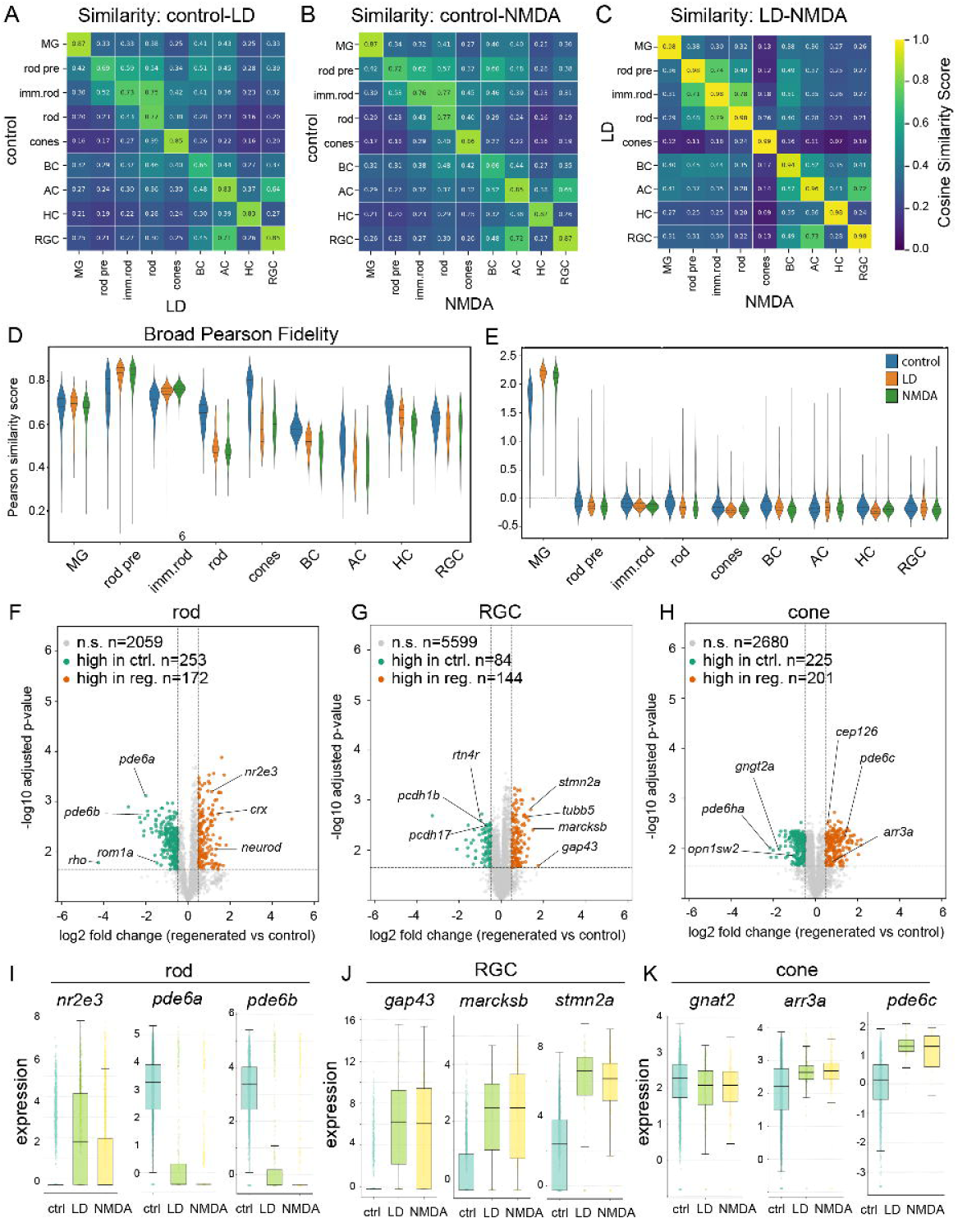
Regenerated neurons transcriptionally resemble endogenous counterparts. (A-C) Cosine similarity analysis between control and light-damaged (A), control and NMDA-injected (B), and light-damaged and NMDA injected (C) samples. (D) Broad Pearson fidelity score across different cell types. (E) Off-target module score across different cell types using Muller glia-specific genes. (F-H) Volcano plots for rods (F), RGCs (G), and cones (H). (I-K) Box plots showing the expression of selected genes in rods (I), RGCs (J), and cones (K). LD: light lesion; NMDA: N-methyl-D-aspartate; MG: Muller glia; rod pre: rod precursor; imm. Rod: postmitotic immature rod: BC: bipolar cell; AC: amacrine cell; HC: horizontal cell; RGC: retinal ganglion cell; ctrl: control.

To further determine the degree to which regenerated neurons transcriptionally resemble endogenous retinal neurons, we next performed fidelity score analysis by comparing each regenerated cell type to its corresponding control counterpart. Consistent with the similarity analysis, fidelity score analysis showed that regenerated cells from both injury models were most highly correlated with their corresponding endogenous cell types (Fig. 3D), further supporting the conclusion that regenerated neurons broadly reestablished cell type-specific transcriptional programs. Next, to determine if regenerated neurons fully lost the molecular program of their parental Müller glia, we performed module score analysis using a Müller glia-specific gene signature. As expected, Müller glia exhibited the highest module score (Fig. 3E), indicating that Müller glia largely regained their Muller glial transcriptional signature after regeneration. In contrast, all regenerated neurons showed minimal enrichment for the Müller glial signature (Fig. 3E), indicating that regenerated neurons lost the parental Müller glial molecular identity during differentiation.

Finally, to define the residual transcriptional differences between regenerated and endogenous neurons, we performed differential gene expression analysis for each cell type (Fig. 3F-H; Table 2). In regenerated rods, genes associated with immature photoreceptor identity, such as *nr2e3*, *crx*, and *neurod*, were enriched, whereas genes involved in mature phototransduction, including *pde6a, ped6b,* and *rho*, were expressed at lower levels than endogenous mature rods (Fig. 3F,I). Likewise, regenerated RGCs exhibited elevated expressions of genes associated with axon growth and cytoskeletal remodeling, such as *stmn2a*, *tubb4*, *gap43*, and *marcksb* (Fig. 3G,J). In regenerated cones, a subset of phototransduction-related genes remained low compared with endogenous cells (Fig. 3H,K). Similarly, regenerated amacrine, bipolar, and horizontal cells exhibited increased expression of genes associated with immature neuronal states, including *hmgb3a* and *hmgb1a* (Fig. S4). Together, these results indicate that 14 days post injury, the residual transcriptional differences between regenerated and endogenous neurons primarily reflect the immature states of regenerated neurons, rather than differences in cell fate specification. Thus, these data indicate that regenerated neurons largely recover cell type-specific transcriptional identities while continuing to undergo maturation.

### 4. Regenerated amacrine cells re-establish subtype diversity and dendritic morphology

In the vertebrate retina, each major cell type encompasses transcriptionally and neurochemically distinct subtypes with specialized physiological roles(Masland 2012a). Because this cellular diversity is fundamental for visual function and acuity, restoring retinal function requires not only regeneration of major cell classes but also subtype-level diversity and circuit-specific features(Masland 2012a; Wässle 2004). While high-throughput transcriptomics has mapped immense cellular complexity in the mammalian retina, identifying dozens of discrete subtypes within each major neuronal class(Yan et al. 2020), the extent to which Müller glia-mediated regeneration can restore this extensive subtype diversity in the adult zebrafish remains poorly characterized. We therefore examined the cell diversity of regenerated neurons, focusing specifically on the subtypes of amacrine cells, bipolar cells and RGCs.

Amacrine cells represent one of the most heterogeneous retinal populations, defined by distinct transcriptional programs, neurotransmitter identities and laminar-specific dendritic architectures (Connaughton et al. 2004; Masland 2012b; Jusuf and Harris 2009). While over 60 transcriptionally distinct amacrine subtypes have been identified in mice, cross species transcriptomic profiling has mapped 42 conserved orthologous types in zebrafish retina (Yan et al. 2020; Tommasini et al. 2026). Our scRNA-Seq subclustering analysis of the amacrine cell population (control: 10,035 cells; light lesion: 236 cells; NMDA: 867 cells) identified 45 transcriptionally distinct clusters (Fig. 4A-C; Fig. S5A-C). Each cluster is defined by unique molecular signatures, representing both known and potentially novel amacrine subtypes (Fig. S5A-C; Table 3). NMDA-injured retinas contained all 45 clusters, whereas light-lesioned retinas exhibited 32 clusters, likely reflecting lower sampling depth (Fig. S5D,E). These data indicate that Müller glia–derived progenitors regenerate a broad spectrum of amacrine subtypes independent of injury paradigm.

**Figure 4.**
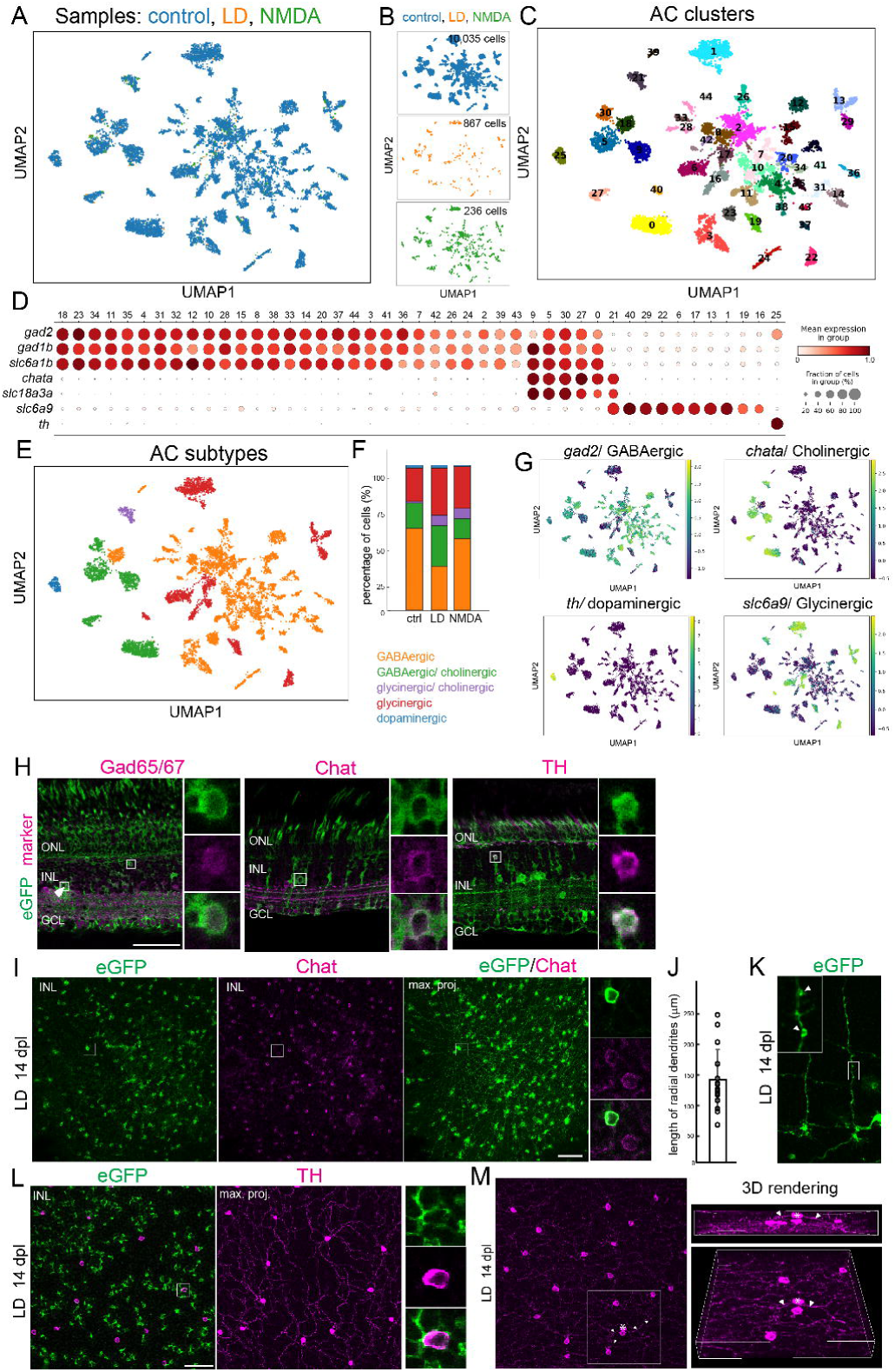
Regeneration restores amacrine subtype diversity and dendritic morphology. (A,B) Integrated UMAP plots showing amacrine cell populations from control, light-damaged and NMDA-injected sample groups. (C) UMAP plot showing regenerated amacrine cell clusters across the three sample groups. (D) Dot plot showing the expression of selected markers for neurotransmitter-defined amacrine cell subtypes (GABAergic: gad2, gad1b, slc6a1b; Cholinergic: chata, slc18a3a; glycinergic: slc6a9; dopaminergic: th). (E) UMAP plot of neurotransmitter-defined amacrine cell subtypes. (F) Proportion of neurotransmitter-defined amacrine subtypes across the three sample groups. (G,H) scRNA-seq mRNA expression of selected marker genes (G) and representative immunostained retinas (H) for GABAergic (gad2/ Gad65/67) Cholinergic (chata/ Chat),dopaminergic (th/ TH) and glycinergic (slc6a9) amacrine subtypes at 14 days after light lesions. (I) Flat-mount retinas immunostained for eGFP and Chat at 14 days post light damage. A single optical section at the inner nuclear layer (left and middle) and Z-stack maximum projection (right) showing Chat+/ eGFP+ regenerated starburst amacrine cells. (J) The length of radial dendrites of regenerated starburst amacrine cells. (K) High magnification image of the dendritic process of regenerated starburst amacrine cells. Arrowheads indicate varicosities. (L) Flat-mount retinas immunostained for eGFP and TH at 14 days post light lesion. A single optical section at the inner nuclear layer (left) and Z-stack maximum projection showing cell bodies of TH+ cells and their dendritic axons within the inner plexiform layer (right). (M) Flat-mount preparation of light-lesioned retina immunostained for TH. Z-stack maximum projection of TH-labeled cells and their dendritic axons. The asterisk indicates apically displaced TH-labeled cells extending dendritic processes (arrows) extending toward the inner plexiform layer. Scale bars: 50 um. LD: light lesion; ONL: outer nuclear layer; INL: inner nuclear layer: GCL ganglion cell layer.

We next classified amacrine clusters based on canonical neurotransmitter markers: GABAergic (*gad2*, *gad1b*, *slc6a1b*), glycinergic (*slc6a9*), cholinergic (*chata*, *slc18a3*), and dopaminergic (*th*) (Fig. 4D). Consistent with previous studies, all identified subtype clusters express markers for the inhibitory transmitters, either GABA or glycine, in mutually exclusive patterns, with 35 identified as GABAergic and 10 as glycinergic (Fig. 4D). Cholinergic clusters co-expressed either GABAergic (5 clusters) or glycinergic (1 cluster) markers (Fig. 4D). Unlike mouse retinas, which possess at least two dopaminergic amacrine subtypes(Yan et al. 2020), only one TH-expressing cluster was identified in zebrafish (Fig. 4D). Notably, all major neurochemical classes were identified in both lesion paradigms (Fig. 4E-G). Immunohistochemistry confirmed the presence of regenerated GABAergic (Gad65/67+), cholinergic (Chat+), and dopaminergic (TH+) amacrine cells in both light-lesioned (Fig.4H) and NMDA-injected retinas (Fig.S5F), validating the transcriptomic classification. Although subtype diversity was broadly restored, relative abundance differed modestly from unlesioned control retinas (Fig. 4F). Regenerated samples exhibited a relative increase in glycinergic (control: 24%, LD: 40%; NMDA: 36%) and cholinergic (control 19%; LD 36%; NMDA 21%) populations and a corresponding decrease in GABAergic cells (control: 74%, LD: 58%; NMDA: 63%), while dopaminergic proportions remained similar (Fig. 4E,F and Fig. S5). Given the limited number of regenerated amacrine cells recovered for scRNA-seq, these proportional shifts should be interpreted cautiously. Nonetheless, these results indicate that regenerated amacrine cells acquire not only appropriate transcriptomic features but also appropriate neurochemical identities, suggesting that the regenerative capacity of the adult zebrafish extends to the production of complex, diverse interneuron subtypes.

Amacrine subtypes are defined not only by transcriptional and neurochemical features but also by highly stereotyped dendritic morphologies and stratification patterns within the IPL(Connaughton et al. 2004; Masland 2012b; Jusuf and Harris 2009). Since cell morphology dictates synaptic connectivity and function, structural recovery of regenerated cells can be utilized as a proxy for the re-establishment of the functional integration of regenerated neurons into synaptic circuits. To examine whether regenerated amacrine subtypes re-establish characteristic dendritic architectures, we focused on two morphologically distinct and well-defined subtypes: cholinergic amacrine cells and dopaminergic amacrine cells, in the dorsal regions of light-lesioned retinas, where eGFP+ cells are sparse (Fig. 1B,C). Cholinergic starburst amacrine cells are characterized by a radially symmetric dendritic tree radiating outward from the central cell body and stratified within the specific sublamina of the IPL(Famiglietti 1983). In the flat-mount retinas, we found abundant eGFP+ cells exhibiting the classical “starburst” morphology, with dendrites radiating from the central soma, and co-labeling with cholinergic marker, Chat, confirmed their identity (Fig. 4I). Their dendritic field diameters range from 63.3um to 245.2um (Ave 136.7±104.1; Fig. 4J). The distal region of dendrite contains numerous varicosities (Fig. 4K), suggesting the formation of functional synapses. These findings indicate that regenerated starburst cells acquire hallmark morphological features.

In the retina, dopaminergic amacrine cells are sparsely distributed but functionally important (Keeley and Reese 2010). Our scRNA-Seq analysis shows that dopaminergic amacrine cells represent about 1% of total amacrine population and constitute a single subtype (Fig. 4E,G). In teleost fish, not mammals, dopaminergic amacrine cells are known as interplexiform cells, extending their process toward the inner plexiform layer and the outer plexiform layer(Dowling and Ehinger 1978; Li and Dowling 2000). Our analysis prioritized processes extending toward the inner plexiform layer, as their dendritic meshwork can be unambiguously traced back to the parent somata. Using TH immunostaining in flat-mount retinas, we identified regenerated dopaminergic neurons that extended fine dendritic processes and integrated into existing circuits within the IPL (Fig. 4L). Although ectopic regenerated dopaminergic amacrine cells were occasionally observed within the outer region of the INL (Fig. 4H, Fig. 4M), their processes also projected toward and integrated within the existing IPL meshwork (Fig. 4M). We observed little evidence of aberrant process targeting, suggesting substantial structural integration of regenerated dopaminergic neurons. Taken together, these data demonstrate that Müller glia–derived progenitors regenerate not only molecularly diverse amacrine subtypes but also re-establish their dendritic architecture and laminar organization. Thus, adult zebrafish retinal regeneration restores interneuron diversity at transcriptional, neurochemical and structural levels, closely recapitulating native populations.

### 5. Regenerated bipolar cells recover ON/OFF diversity and retinal lamination

Retinal bipolar cells are interneurons that receive synaptic input from photoreceptors and transfer visual signals to amacrine and ganglion cells within the inner plexiform layer. They are classified into distinct ON and OFF subtypes that stratify within specific sublayers of the inner plexiform layer. ScRNA-Seq subclustering analysis of the pooled bipolar cells (control: 58,749 cells; LD: 545 cells; NMDA 844 cells) identified 27 transcriptionally distinct subtypes, each defined by a unique molecular signature (Fig. 5A,B; Fig. S6A,B; Table 4). Although the regenerated samples contained all bipolar cell subtypes (Fig. 5A,B), they showed dramatic change in the distribution of cell fraction in each cluster compared with control (Fig. 5C). Clusters 19, 15, 26, 20,18 and 14 displayed increases, whereas clusters 0, 1, 2, 3 and 8 significantly decreased in the regenerated samples relative to control (Fig. 5D). To characterize the molecular features of the altered bipolar cell populations, we performed differential gene expression analysis. Clusters 19, 15 and 26 were enriched for genes associated with late-stage progenitors and immature neurons (*nfia*, *nfib*, *nfixa*, *nifxb*, *hmgb1a*, *hbgm1b* and *hbgm3a*) as well as genes involved in structural remodeling (*stmn1b*, *rtn4a*, *rtn4r* and *marcks1a*). Consistent with these findings, module score analysis showed elevated expression of immature neuronal gene signatures in these clusters (Fig. 5E; Fig. S6C,D), indicating these expanded clusters represent immature bipolar cells undergoing differentiation. To establish whether regenerated bipolar cells reestablished ON and OFF subtype identities, we classified clusters using established ON (*grm6a*, *trpm1b*, *prkcaa, trpm1a, and grm6b*) and OFF (*grik1b*, *neto1*, *neto1l* and *grik4*) marker genes. This analysis identified 11 ON and 13 OFF bipolar cell subtypes (Fig. 5F,G). Both ON and OFF populations were present in regenerated retina, and this was validated by immunostaining for ON bipolar marker, anti-aPkc and ON/OFF marker, anti-Calb2 (Fig. 5H). Notably, the two most consistently expanded mature bipolar populations, clusters 15 and 18, corresponded to OFF bipolar cell subtypes characterized by enriched expression of neto1l and grik4, respectively (Fig. 5C,D,F). These results indicate that regeneration restores both ON and OFF bipolar cells. However, the relative abundance of individual bipolar cell subtypes is quantitatively altered.

**Figure 5.**
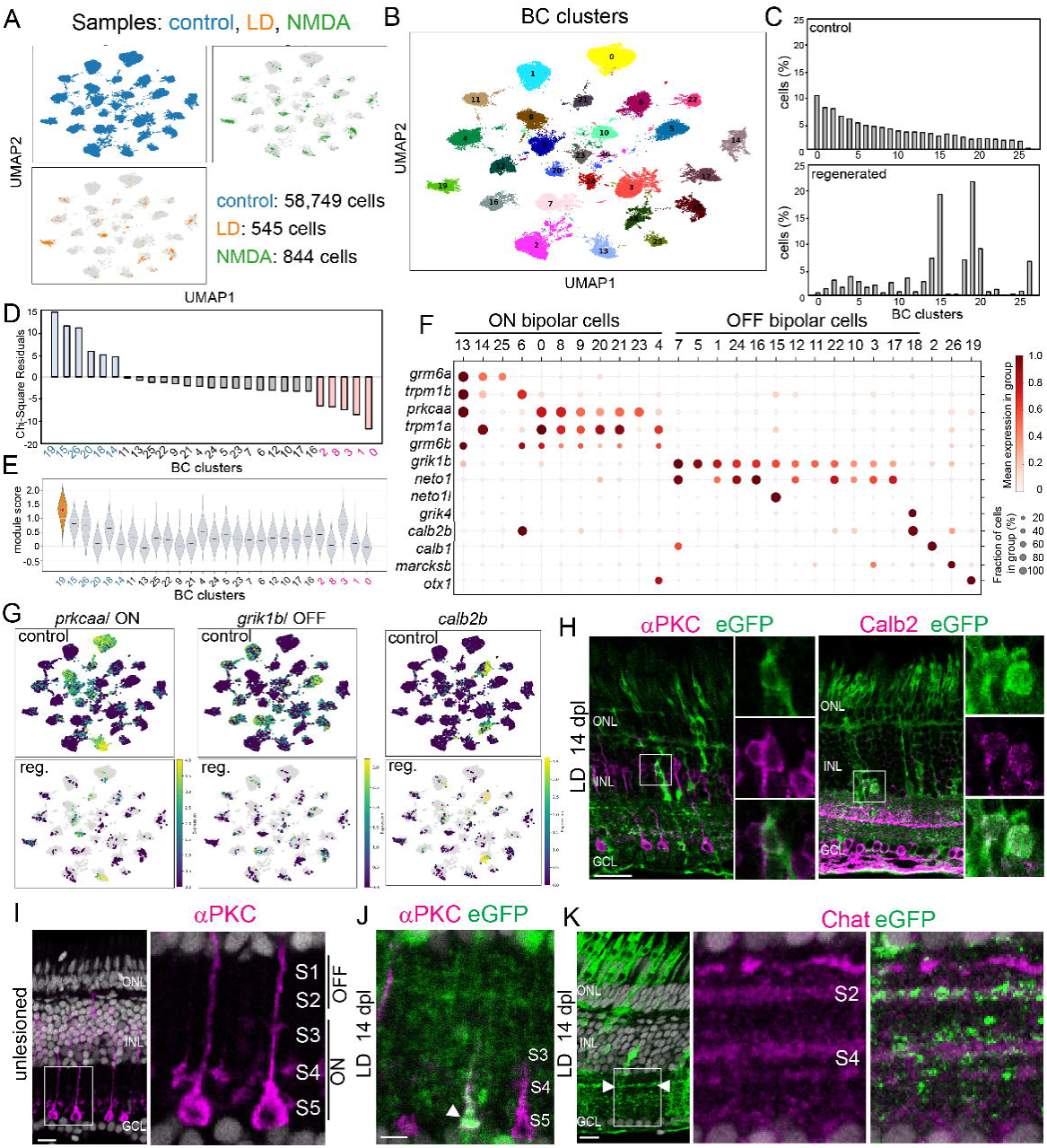
Regenerated bipolar cells show subtype diversity and cell-type specific laminar targeting. (A) Integrated UMAP plot of retinal bipolar cell populations from control, light-damaged, and NMDA-injected sample groups. (B) UMAP plot showing bipolar cell clusters across the three sample groups. (C) Proportion of bipolar cell subtypes in control (top) and regenerated (bottom) retinal samples. (D) Pearson’s chi-square test showing the composition shift following regeneration. (E) Immature gene module scores across bipolar cell subtypes. (F) Dot plot showing the expression of selected genes for ON and OFF bipolar cells. (G) UMAP plots of selected marker genes. (H) Representative immunostaining for aPKC and Calb at 14 days post light lesion. (I-J) Representative immunostaining for aPKC in unlesioned (I) and 14 days post light-lesioned (J) retinas. (K) Representative immunostaining for Chat at 14 days post light lesion. Scale bars: H 50 um; I-K 20 um. LD: light lesion; ONL: outer nuclear layer; INL: inner nuclear layer: GCL ganglion cell layer; S: sublamina.

The inner plexiform layer (IPL) is stratified into five sublaminae (S1 to S5), which can be divided into two main functional halves, the OFF pathway (outer IPL, S1 and S2) and the ON pathway (inner IPL, also called sublamina b, S3, S4, S5) (Fig. 5I). To determine if regenerated neurons conform to these precise sublaminae within the IPL, we used established markers that define individual IPL sublaminae. In the control retinas, the aPKC antibody labels the axon terminals of ON bipolar cells within S3, S4, and S5 (Fig. 5I). In the regenerated retinas, the axons of aPKC+ bipolar cells terminated deep within the ON sublaminae, integrated among the terminals of the endogenous cells (Fig. 5J; Fig. S6E). To assess laminar targeting within the amacrine cell network, we used Chat immunostaining, which labels S2 and S4 sublaminae (Fig. 5K). A prominent eGFP-labeled band precisely colocalized with Chat+ immunostaining signal (Fig. 5K; Fig. S6F). This proper spatial stratification pattern of regenerated neurons demonstrates that regenerated retinal neurons accurately restore cell-type specific laminar targeting within the IPL.

### 6. Regenerated RGCs recapitulate molecular diversity and undergo microglia-mediated refinement

RGCs comprise a highly heterogeneous population of projection neurons. To determine whether different RGC subtypes are regenerated following retinal injury, we performed scRNA-Seq subclustering analysis of the pooled RGC datasets (control: 4,000 cells, LD: 150 cells, NMDA 430 cells). Our analysis identified 37 transcriptionally distinct clusters, each defined by a unique molecular signature (Fig. 6A-C; Fig. S7A,B; Table 5). Regenerated RGCs populated 37 clusters (Fig. 6D,E), including *atoh7*+ immature RGCs (Fig. S7A) and *eomesa*+ intrinsically photosensitive RGCs (Fig. S7B). Although the scRNA-seq analysis showed that regenerated RGCs exhibited transcriptomic profiles highly similar to their endogenous counterparts, we observed that a significant number of RGCs were ectopically located (Fig.6F-I; Fig. S7C-F). In unlesioned control retina, RGC somata were strictly confined to the GCL (Fig. S7C,D). In contrast, 43.6% and 32.7% of eGFP+ regenerated RGCs were mislocalized in light-lesioned and NMDA-injected retinas, respectively (Fig. 6F-I; Fig. S7D-F). These findings indicate that RGC subtype specification is established relatively early during regeneration, whereas accurate laminar positioning remains incomplete. We next asked whether displaced RGCs are selectively eliminated over time. Quantification showed that regenerated RGC numbers declined significantly between 14 and 28 days after both light lesion and NMDA injury, with comparable reductions observed in both the GCL and ectopic locations (Fig. 6J). This concurrent decline for RGCs across all layers indicates a broad elimination process affecting regenerated RGCs regardless of their position.

**Figure 6.**
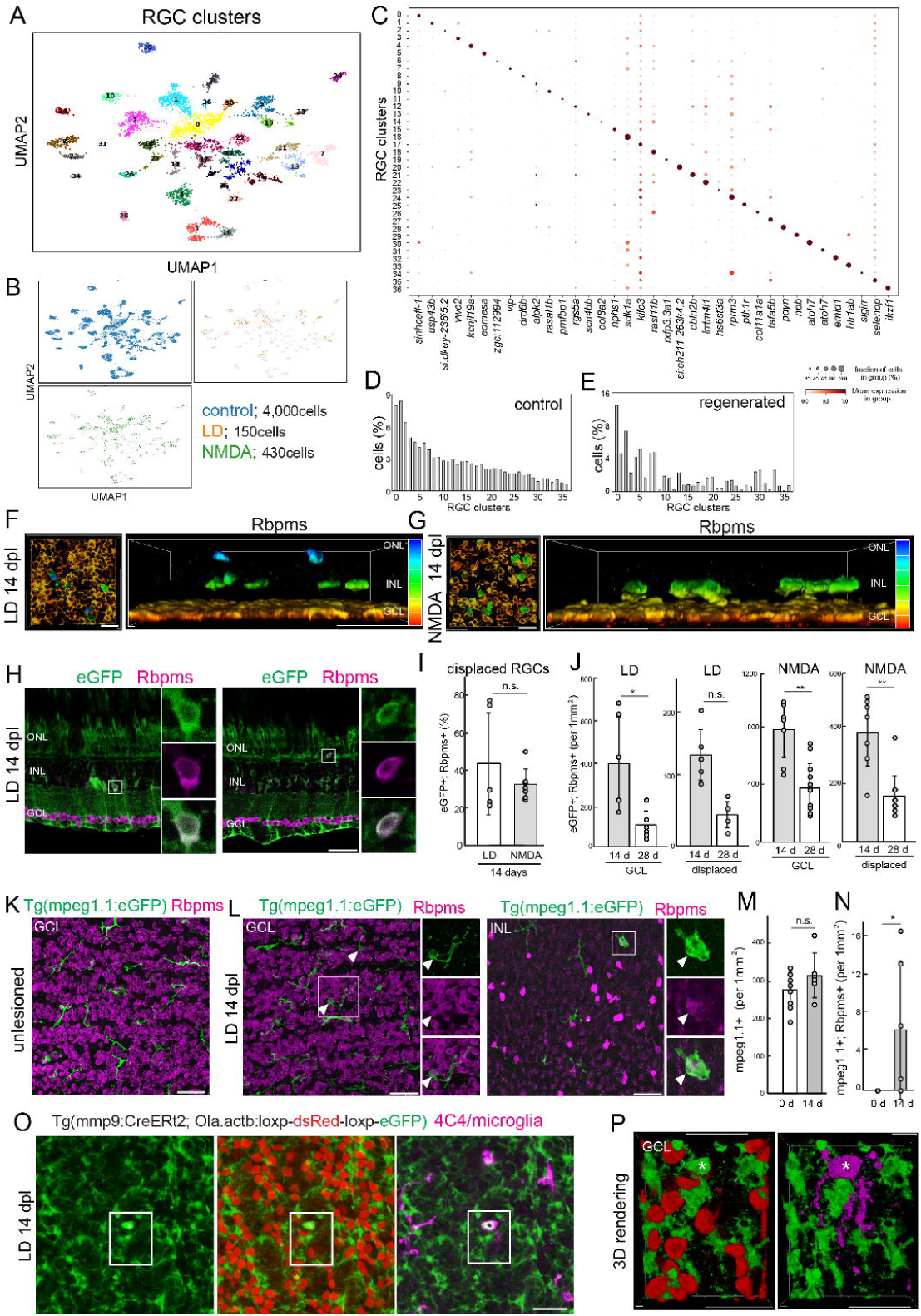
Regenerated RGCs exhibit subtype diversity, ectopic positioning, and microglia-mediated elimination. (A,B) UMAP plot of RGC populations from control, light-lesioned, and NDMA injected sample groups. (C) Dot plot showing the expression of selected markers for RGC clusters. (D,E) Proportion of RGC clusters in control (D) and regenerated (E) samples. (F,G) Flat-mount retina following light lesion (F) and NMDA injury (G) immunostained for Rbpms. Maximum projection of Z-stack series with depth color-coded (left) and cross section view of 3D rendering with depth color-coded (right). (H) Retinal cross section immunostained for Rbpms at 14 days post light lesion. (I) The percentage of displaced eGFP+/Rbpms+ cells among total eGFP+/Rbpms+ cells. (p=0.61) (J) The number of eGFP+/Rbpms+ cells in the RGC layer and displaced populations at 14 and 28 days following light lesion (GCL: p=0.02; displaced: p=0.08) and NMDA injury (GCL: p=0.0005; displaced: p=0.0001). (K,L) Flat-mount retinal preparation of microglia reporter, Tg(mpeg1.1: eGFP), immunostained for Rbpms in control (J) and 14 days post light lesion (K) in the GCL and INL. Box region and arrowhead indicate microglia phagocytosing Rbpms+ cells. (M) The number of Tg(mpeg1.1:eGFP)+ microglia in the ganglion cell layer in unlesioned and at 14 days post light lesion (p=0.0932). (N) The number of Tg(mpeg1.1:eGFP)+/ Rbpms+ cells (p=0.0133). (O) Flat-mount Tg(mmp9:creERT2; Olactb:loxp-dsRed-loxp-eGFP) retina immunostained for eGFP and 4C4 at 14 days post light lesion. (P) 3D rendering of the boxed region in O. Asterisk indicates eGFP+/4C4+ cell. Scale bars: F,G,O,P 20um; H.K,L 50 um. LD: light lesion; dpl: days post lesion; NMDA: N-methyl-D-aspartate; dpi: days post injection, GCL ganglion cell layer; INL: inner retinal layer; OPL: outer plexiform layer: ONL: outer nuclear layer.

During retinal development, nascent RGCs are initially over-produced and subsequently eliminated in an activity-dependent manner(Guerin et al. 2006; Cellerino et al. 2000). During this elimination process, microglia, the innate immune cells of the central nervous system, play a crucial role(Anderson et al. 2019). We therefore investigated whether microglia act similarly to cull regenerated RGCs. Using the transgenic microglial reporter line, *Tg(mpeg1.1: eGFP)*, we observed eGFP+ microglia actively phagocytosing Rbpms+ RGCs in both the GCL and INL at 14 days post injury (Fig. 6K-N). Furthermore, our lineage tracing demonstrated that microglia phagocytosed eGFP+ regenerated neurons within the GCL (Fig. 6 O,P). Taken together, these results suggest that regenerated RGCs are initially overproduced and subsequently refined by microglia-mediated pruning, resembling the refinement that occurs during development.

### 7. Regenerated RGCs re-establish long-range retinotectal projections

We next examined whether regenerated RGCs successfully re-establish long-range axonal projection to their appropriate central targets. Unlike mammals, zebrafish RGC axons project exclusively through the optic chiasm to innervate the contralateral optic tectum (Fig. 7A). To visualize regenerated axons, we injected NMDA into the right eye and processed whole brains for light-sheet microscopy and flat-mount optic tecta for confocal microscopy. As expected, NMDA injury induced eGFP exclusively in the damaged retina (Fig. 7B). By 7 days post injury, regenerated eGFP+ axons reached the contralateral tectum and were observed along its peripheral margins (Fig, 7D,D’). By 14 days after injury, regenerated axons formed prominent fasciculated bundles extending along the anterior-posterior axis of the tectum (Fig. 7E,E’; Fig. S8A). Cross sections of the tectum further demonstrated that regenerated RGC axons accurately innervated the stratum opticum (Fig.7F), faithfully recapitulating the projection pattern observed in uninjured animals following anterograde labeling with cholera toxin subunit B (Fig. 7G). Flat-mounted ipsilateral tecta (Fig. S8B) and light-sheet imaging of whole brains (Fig. 7C) showed negligible aberrant RGC projections. Similar patterns of tectal reinnervation were observed following light-induced retinal injury (Fig. S8C,D), suggesting that the reinnervation of RGC axons into the optic tectum does not strictly require the loss of RGCs. Taken together, these results demonstrate that regenerated RGC axons accurately reinnervate their appropriate target within the contralateral optic tectum.

**Figure 7.**
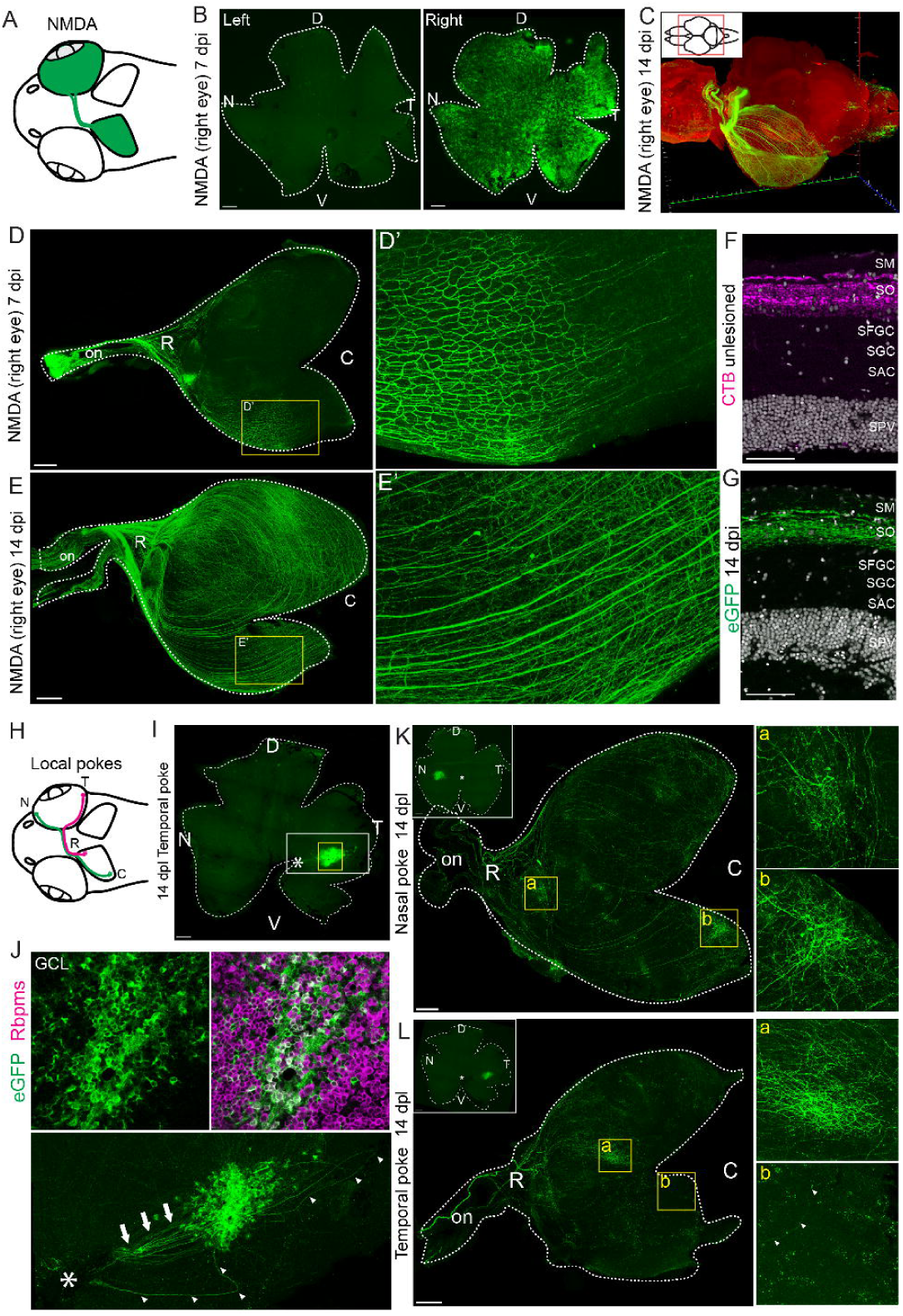
Regenerated RGC axons re-establish retinotectal projections. (A) Schematic of contralateral projections of retinal ganglion cells in the zebrafish visual system. (B) Flat-mount retina at 7 days following NMDA injection into the right eye. (C) 3D rendering of the whole cleared brain at 14 days following NMDA injection into the right eye, imaged by light-sheet microscopy. (D, E) Flat-mount preparation of contralateral tectum immunostained for eGFP at 7 days (D, D’) and 14 days (E, E’) following NMDA injection into the right eye. (F) Cross section of the tectum from unlesioned control fish 2 days after CTB injection. (G) Cross section of tectum at 14 days post NMDA lesion. (H) Schematic of retinotectal topography in the zebrafish visual system, showing nasal RGC axons projecting to the caudal tectum and temporal RGC axons projecting to the rostral tectum. (I) Flat-mount retinal preparation at 14 days post a poke injury to the temporal region of the retina. Asterisk indicates optic disc. (J) High magnification image of boxed regions immunostained for Rbpms. Asterisk and arrows indicate optic disc and axons, respectively. Arrowheads indicate misrouting axons. (K,L) Flat-mount preparation of contralateral tectum immunostained for eGFP at 14 days following nasal (K) or temporal (L) poke lesions. Scale bars: B, D, E, I, K, L 200um; F, G 50um. D: dorsal; V: ventral; N: nasal; T: temporal; op: optic nerve; R; rostral; C:Caudal; SM: Stratum Marginale; SO: Stratum Opticum; SFGC: Stratum Fibrosum et Greseum Superficale; SAC: Stratum Griseum Centrale; SPV: Stratum Album Centrale; GCL ganglion cell layer..

During normal visual system development, RGC axons form a highly ordered retinotopic map within the optic tectum (Kita et al. 2015). To determine whether regenerated RGCs also restore this topographic organization, we performed localized puncture lesions in either the nasal or temporal retina. This approach takes advantage of the native retinotectal map, in which temporal RGCs project to the rostral tectum whereas nasal RGCs innervate the caudal tectum (Fig. 7H). Localized injury resulted in the generation of eGFP+ RGCs that were confined to the injury site, as confirmed by eGFP+/Rbpms co-labeling (Fig.7I,J). Although most regenerated RGC axons extended directly toward the optic disc, some initially followed aberrant trajectories before undergoing course correction and redirecting toward the optic nerve head (Fig. 7J, arrowheads), suggesting that regenerated axons may respond to local guidance cues during intraretinal axon extension. By 14 days post injury, regenerated RGC axons arising from nasal retinal lesions projected predominantly to the caudal tectum (Fig. 7K), whereas regenerated axons from temporal lesions selectively innervated the rostral tectum (Fig. 7L). Although occasional eGFP+ axons from temporal lesions extended into the caudal tectum, these axons failed to branch or elaborate terminal arbors (Fig. 7L). These results demonstrate that regenerated RGC axons not only re-establish long-range connectivity but also accurately reconstruct retinotectal topography.

## Discussion

A major challenge in regenerative neuroscience is not only replacing lost neurons, but also rebuilding appropriate cellular diversity and circuit organization required for normal function. During development, the vertebrate retina is formed through a highly coordinated sequence of cell fate specification, migration, laminar organization, and circuit formation. Whether adult regeneration can faithfully reconstruct this complex organization has remained a fundamental question in the field. By combining lineage tracing, single-cell transcriptomics and structural analyses, we demonstrated that Müller glia–mediated regeneration reconstructs the adult zebrafish retina across multiple scales. Regenerated neurons recover endogenous molecular identities, re-establish neuronal subtype diversity, restore characteristic tissue organization, and rebuild long-range retinotectal projections. Together, these findings establish the adult zebrafish retina as a powerful model for uncovering the molecular and cellular mechanisms that enable faithful neural circuit reconstruction.

Our data show that Müller glia-derived progenitors regenerate all major retinal cell types following either injury paradigms, while the relative abundance of regenerated neurons is biased toward the pattern of neuronal loss. These findings support a model in which retinal regeneration is governed by the interplay between conserved intrinsic neurogenic programs and extrinsic injury-induced signals. In this model, intrinsic neurogenic programs enable Müller glia-derived progenitors to generate multiple retinal lineages, whereas extrinsic injury-induced signals quantitatively bias regenerative output toward the neuronal populations most affected by damage. This interpretation is supported by previous studies demonstrating a shared core regenerative program activated in Muller glia across different injury paradigms (Hoang et al. 2020); (Lyu et al. 2023; Emmerich et al. 2023, 2024). Likewise, the conserved temporal expression of developmental competence factors in Muller glia-derived progenitors (Lahne, Brecker, et al. 2020) further supports that regeneration largely recapitulates intrinsic developmental programs. How injury-specific signals shape the magnitude and composition of regenerative output, however, remains incompletely understood. We previously identified the extracellular matrix protease Mmp9 as an injury-induced regulator of Müller glia-mediated regeneration (Lyu et al. 2023). Furthermore, both our work and others have shown that inflammatory signaling influences the proliferation of Müller glia-derived progenitors and their fate specification(Silva et al. 2020; Nagashima and Hitchcock 2021; Magner et al. 2022). Determining whether different lesion paradigms elicit distinct inflammatory or extracellular signaling environments that interact with intrinsic programs to direct specific lineages will be an important direction for future studies.

An important unanswered question is how regenerative output is matched to tissue demand. Our study, together with prior work(Powell et al. 2016; Yoshimatsu et al. 2016; Lyu et al. 2023), demonstrates that Muller glia-derived progenitors regenerate a broad repertoire of neuronal types, regardless of whether the corresponding endogenous cell type was directly lost. A similar phenomenon has been reported in other regenerative contexts, including zebrafish spinal cord (Reimer et al. 2008), suggesting that broad neuronal production may represent an evolutionarily conserved strategy favoring robust neuronal replacement followed by refinement through survival, maturation, and circuit integration. Consistent with this model, we found that a subset of regenerated RGCs was removed by microglial phagocytosis, reminiscent of developmental processes in which excess neurons and synaptic connections are selectively pruned to optimize neural circuit assembly (Brown and Neher 2014; Mosser et al. 2017; Faust et al. 2021). Whether a similar refinement process is required for full functional recovery after injury, however, remains unclear. Unlike the developing nervous system, the adult regenerative environment may tolerate additional or ectopically positioned neurons if they can establish appropriate functional connections. It remains unknown whether microglia selectively eliminate regenerated neurons based on their maturation state, neuronal activity, synaptic integration, or competition with neighboring neurons. Defining how regenerative systems balance neuronal production with selective refinement will be critical for developing regenerative strategies that achieve precise neuronal replacement and functional circuit reconstruction.

Our findings demonstrate that Müller glia–mediated regeneration reconstructs the adult zebrafish retina across multiple levels, from molecular identity to local and long-range neural circuits. Across all major retinal classes, regenerated neurons closely match their endogenous counterparts at the molecular level, with the remaining transcriptional differences largely reflecting ongoing maturation. At the circuit level, regenerated bipolar and amacrine cells re-established their morphologies and laminar organization, while regenerated RGCs reinnervated the contralateral optic tectum and largely restored retinotopic organization following focal retinal injury. These findings extend previous optic nerve regeneration studies (Becker and Becker 2007; Bollaerts et al. 2018) by demonstrating that newly generated RGCs can also navigate to appropriate targets and reconstruct visual projections. Collectively, our results suggest that the adult zebrafish retina and brain retain instructive positional and guidance cues that enable accurate reconstruction of neural circuits long after development is complete. Future longitudinal molecular, physiological, and behavioral studies will be needed to determine when regenerated neurons achieve full functional maturity.

Several limitations should be considered when interpreting our findings. First, most molecular and structural analyses were performed at a relatively early stage of regeneration (14 days after injury), before regenerated neurons are expected to achieve full physiological maturity. Future studies using electrophysiological and behavioral assays will be needed to determine whether visual function is fully restored. Second, the number of lineage-traced cells recovered by scRNA-Seq, particularly for several neuronal subtypes, was relatively modest, limiting the statistical power to detect subtle differences in subtype composition or transcriptional states.Third, while we performed detailed subtype analysis of amacrine cells, bipolar cells and RGCs, comparable analyses of photoreceptors and horizontal cells would be needed to fully assess the diversity of regenerated neurons. Finally, because our study focused on acute injury paradigms, the extent to which these findings generalize to chronic retinal degenerative diseases remains an important question for future investigation.

In summary, our study demonstrates that Müller glia-mediated regeneration reconstructs the adult zebrafish retina across multiple biological scales, from molecular identity and neuronal subtype diversity to neural circuitry. Although spontaneous retinal regeneration is extremely limited in mammals, recent studies from our group and others have shown that Müller glia-derived neurons generated in the adult mouse retina can respond to light stimulation (Jorstad et al. 2017; Todd et al. 2022; Yang et al. 2025; Hoang et al. 2020; Le et al. 2024), indicating that the adult mammalian retina retains a degree of regenerative plasticity. Our findings suggest that successful neuronal replacement requires coordinated restoration of neuronal identity, cellular organization and circuit connectivity. Defining the molecular and cellular mechanisms that enable high levels of regenerative fidelity in the adult zebrafish retina will provide a foundation for developing regenerative therapies for the injured mammalian retina and, more broadly, the adult central nervous system.

## Author contribution

Conceptualization: M.N, P.H and T.H. Methodology: M.N, L.J, L.R.S, S.A, Z.F, P.H, and T.H. Investigation: M.N, L.J, L.R.S, S.A, Z.F, P.H, and T.H. Formal Analysis: M.N, L.J, L.R.S, S.A, Z.F, P.H, and T.H. Supervision: MN, PH and T.H. Funding acquisition: P.H and T.H. Writing—original draft: M.N and T.H. Writing—review and editing: M.N, L.J, L.R.S, S.A, Z.F, P.H, and T.H

## Acknowledgement

This work was supported by NIH P30EY007003 Vision Research Core grant and Research to Prevent Blindness unrestricted Grant to the Department of Ophthalmology, and NIHR21EY034182 grant to P.H.

## Resource availability

### Lead contact

Further information and requests for resources and reagents should be directed to and will be fulfilled by the lead contact

### Data and code availability

- Raw scRNA-seq data have been deposited at the Gene Expression Omnibus (GEO) and are publicly available as of the date of publication. Accession number: GSE319801.
- Original code for this study is available at: https://github.com/SherineAwad/Zebrafish_scRNASeq_scanpy
- Interactive companion site is available at https://kec.singlecell.med.umich.edu/.
- Any additional information required to reanalyze the data reported in this paper is available from the lead contact upon request

## Experimental model and subject details

### Zebrafish

Wild-type, AB strain zebrafish (Danio rerio; ZIRC, University of Oregon, Eugene, OR), lineage tracing reporter line, *Tg(mmp9:creERt2; Olactb:loxP-dsred2-loxp-egfp)*(Ando et al. 2017), microglial reporter line *Tg(mpeg1.1:eGFP)* (Ellett et al. 2011) were maintained at 28C on a 14/10 h/ light/ dark cycle with standard husbandry procedures. Adults were of either sex and used between 4-10 months of age. To generate lineage tracing lines, male *Tg(mmp9:creERt2)tyt208*(Ando et al. 2017) were out crossed with female *Tg(Olactb:loxP-dsred2-loxP-egfp)tyt201Tg*(Yoshinari et al. 2012). Male Cre-drivers were used to avoid premature recombination in the germline or early embryonic stage due to maternal deposition of Cre mRNA/protein. All experimental protocols were approved by the University of Michigan Institutional Animal Care and Use Committee.

## Method details

### Pharmacological treatment

To induce Cre-mediated recombination, zebrafish were treated with 4-hydroxytamoxifen (4-OHT) via immersion. A stock solution of 2.5 mM 4-OHT was prepared in Ethanol and stored at -20C. For treatment, the stock was diluted in system water to a final concentration of 2.5uM. Adult fish were subjected to four consecutive, 12 hours of overnight treatments(Pinzon-Olejua et al. 2017). In between each treatment, animals were placed in a system of water. To prevent photodegradation of the Tamoxifen, tanks were kept in the dark during the incubation periods. To assay cell proliferation, EdU (5-Ethynyl-2’-deoxyuridine, Thermo Fisher Scientific) was administered intraperitoneally. Fish were anesthetized using pharmaceutical grade/FDA approved MS-222/ Tricaine S (0.02%/ 20 mg/ 100ml, Western Chemical Inc., Ferndale, WA) dissolved in system water containing 0.02% sodium bicarbonate. 20 uL of 1mg/ml EdU in PBS was injected intraperitoneally using a 30Gx1/2 needle with 1ml syringe at 24, 48, 72, and 96 hours post lesion.

### Retinal lesions

To selectively damage photoreceptors, an intense light lesion was used(Bernardos et al. 2007; Taylor et al. 2012). Zebrafish were exposed to 100,000 lux light from an EXFO X-Cite 120 W metal halide lamp for 30 min. To damage inner retinal neurons, 0.5ul of 100 mM NMDA in PBS was injected intravitreally. Using a 30Gx1/2 needle, a small incision was made in the cornea. A hamilton syringe equipped with a blunt 30G needle was carefully inserted behind the lens, then 0.3 ul of NMDA solution was delivered to the vitreous. Local poke lesion was performed as described previously (Senut et al. 2004) by inserting a 30Gx1/2 needle into the nasal or temporal aspect of the eyes through the sclera. NMDA injections and local poke lesions were performed under anesthetized conditions using 0.02% MS-222.

### Fixation and tissue processing

Following anesthesia with 0.02% MS-222, eyes were enucleated and placed in 4% paraformaldehyde in 0.1M phosphate buffer (pH7.4) containing 5% sucrose overnight at 4C. To prepare tectal tissues, zebrafish heads were placed in fixative overnight at 4C. After rinsing with 5% sucrose solution, brain tissues were carefully dissected in PBS. Tissues were cryoprotected with 20% sucrose solution, then embedded in OCT/ 20% sucrose solution. Retinal and tectal samples were sectioned at 6 um using Leica CM3050S cryostat.

### Immunohistochemistry and imaging

Immunohistochemistry was performed as described in our previous studies(Nagashima et al. 2020; Silva et al. 2020). Briefly, sections were blocked in blocking reagents containing 20% normal goat serum/0.5% Triton X-100 in pH7.4, with 0.1% sodium azide. Primary antibodies were diluted in a dilution buffer containing 1% normal goat serum/0.5% Triton X-100 in pH7.4, with 0.1% sodium azide and incubated 4C overnight. AlexaFluor secondary antibody (Thermo Fisher Scientific) incubation was performed at room temperature for 2 hours. Slides were stained with Hoechst 33342 (Thermo Fisher Scientific) for nuclear staining. Sections were mounted with ProLong Gold (Thermo Fisher Scientific). For goat anti-Choline acetyltransferase antibody, horse serum was utilized as the blocking and dilution agents in place of goat serum to prevent cross-reactivity. Flat-mount immunocytochemistry was performed as previously described(Silva et al. 2020; Nagashima et al. 2020) using blocking (10% normal goat serum/1% Tween 20/1% Triton X-100/1% DMSO in PBS, pH 7.4, with 0.1% sodium azide) and dilution (0.5% normal goat serum/1% Tween 20/1% Triton X-100/1% DMSO in PBS, pH 7.4, with 0.1% sodium azide) reagents. To prepare tectal samples for lightsheet microscope imaging, whole-brains were immunostained as described above, optical clearing was performed using EasyIndex solution. Tissues were incubated in a 50%EasyIndex solution (diluted in distilled water for 3 hours at room temperature. Brains were then transferred to 100% EasyIndex and incubated overnight to complete optical transparency. For subsequent dissection, tissues were rehydrated via incubation in PBS. Optic tectal tissues were then carefully dissected and prepared for flat-mount imaging.

### EdU and TUNEL assays

Visualization of EdU labeled cells and TUNEL staining were performed using Click-iT EdU Alexa Fluor 647 Imaging kit (Thermo Fisher Scientific), and In Situ Cell Death Detection Kit, TMR red (Millipore Sigma), respectively, according to the manufacturer’s instructions. For TUNEL staining, slides were treated with PBS containing 1% sodium citrate/ 1% Triton X-100 at 4C. After rinsing with PBS, slides were incubated with a TUNEL reaction cocktail for 30 min at 37C. For EdU detection, freshly prepared Click-iT reaction cocktails were treated for 30 minutes at room temperature.

### Imaging and analysis

Fluorescence images of retinal sections were captured using Leica Stellaris 8 Falcon Confocal Microscope. Flat-mount retinal preparations were imaged using Leica SP5 Confocal Microscope or Leica Stellaris 8 Falcon Confocal Microscope. Light-sheet imaging was captured using Zeiss Lightsheet 7 following overnight tissue clearing treatment with EazyIndex. Cell counts were performed on Leica Application Suite X (Leica Microsystems). For retinal cross-section, cell counts were quantified across a 300 um linear length of the retina. To ensure representative sampling, 2-3 non-adjacent sections were analyzed per retina. The resulting counts were averaged per individual and normalized. For flat-mount preparation, cells in 22,500mm2 were quantified and normalized. To statistically compare cell counts between two groups, an independent t-test was performed. Statistical analysis across time points was performed using a one=way Analysis of Variance (ANOVA).

### Retinal dissociation, FACS isolation and scRNA-Seq

Retinas from 4 adult fish were dissected in ice-cold Hibernate-A (Gibco A1247501) and dissociated using the Papain Dissociation System (LK003150, Worthington) with the following modifications. Retinas were incubated in papain enzyme buffer at 4C for 20min, then 28C for 30 min, with tube mixing by inversion every 5 min. The tissue was centrifuged at 200xg for 2 min and papain solution removed, then dissociated in ice-cold Hibernate-A buffer by trituration. The single cell suspension was subjected to gradient centrifugation through ovomucoid albumin at 200xg for 7min to exclude dead cells. Live pelleted cells were resuspended in HBAG buffer containing Hibernate-A, B-27 supplement (Thermo Fisher 17504044), and GlutaMAX (Thermo Fisher 35050061), filtered through 50um filter, and then subjected to FACS using a Sony MA900 cell sorter for GFP+ cells into HBAG buffer. Sorted cells were mixed with BSA at 0.3% final concentration to reduce potential cell clumping before centrifugation at 400xg for 5min to concentrate, then resuspended in a desired volume of HBAG buffer to reach a concentration of 500-1500 cells/uL. Cell viability and counts were determined via trypan blue staining and haemocytometer. Cells were submitted to the University of Michigan Advanced Genomics Core for 3’ scRNA-seq. GFP+ cells (∼15k) were loaded into the 10X Genomics Chromium Single Cell System and libraries were generated using V4 chemistry following the manufacturer’s instructions. Libraries were sequenced on the Illumina NovaSeq platform (500 million reads per library).

### scRNA-Seq data processing and analysis

Single-cell RNA-sequencing data were analyzed using the Scanpy framework (v1.9.3) together with AnnData (v0.12.8). Quality control filtering was applied to remove low-quality cells and potential multiplets. Cells were retained if they expressed more than 800 and fewer than 6,000 genes, contained between 1,200 and 30,000 total transcripts, and exhibited less than 25% mitochondrial gene expression. In addition, cells expressing fewer than 100 genes were excluded, and genes detected in fewer than three cells were removed from further analysis. Following quality control, gene expression values were normalized to account for differences in sequencing depth and log-transformed. Highly variable genes were identified and used for downstream analyses. Dimensionality reduction was performed using principal component analysis to capture the major sources of transcriptional variation. To correct for batch effects between samples, data integration was performed using the Harmony algorithm. Batch-corrected principal components were used to construct a nearest-neighbor graph, which served as the basis for downstream analyses. Cell populations were identified using graph-based clustering with the Leiden algorithm.

Following initial clustering and cell type annotation, specific cell populations were subset based on annotated cell type identities. Cells belonging to selected cell types were extracted from the integrated dataset and re-analyzed independently to enable higher-resolution characterization of within-population heterogeneity. Differential gene expression analysis was performed using a nonparametric Wilcoxon rank-sum test as implemented in Scanpy. Depending on the analysis context, differential expression was assessed either between annotated cell clusters or between specified sample groups. For cluster-level analyses, genes were ranked for each cluster relative to all remaining cells, whereas sample-level comparisons were performed between cells derived from two selected samples. For each comparison, gene-level statistics including Wilcoxon scores, log fold changes, and multiple-testing–adjusted p-values were calculated. Differentially expressed genes were ranked based on statistical significance or effect size and used for downstream visualization and interpretation.

### Cell-type similarity and composition analyses

Cell-type similarity was evaluated using pseudobulk centroid-based cosine similarity. Cells were grouped by annotated cell type and condition, and mean expression profiles were calculated for each cell type within Control, LD, and NMDA using raw.X expression values. For the heatmap analysis, highly variable genes were selected across retinal cells, and the final heatmaps used the top 2,000 HVGs (hvg_2000). Pairwise cosine similarities were then calculated between reference and query cell-type centroids for Control versus LD, Control versus NMDA, and LD versus NMDA comparisons.

To remove a Rod outlier population identified by UMAP/Leiden analysis, Rod cells assigned to Leiden clusters 30 and 86 were excluded in memory before calculating centroids; the original h5ad file was not modified. Heatmaps were ordered as MG, MGPC, PR precursors, Rod, Cones, BC, AC, HC, and RGC.

Fidelity score analysis was performed using a Pearson correlation score comparing each cell to its corresponding control cell-type centroid. Control-derived marker genes were selected for each cell type using differential enrichment criteria of avg_log2FC >= 0, pct_in_target >= 0.05, adjusted p <= 0.05, no pct_in_other cutoff, and a maximum of 2,000 genes per cell type. Fidelity scores were calculated for each cell type. Higher fidelity scores indicate stronger similarity to the matching control cell-type transcriptional profile. Müller glia off-target score was quantified using a module score based on the top 100 Müller glia-enriched genes. The gene set was selected from the Müller glia differential enrichment ranking and scored using scanpy.tl.score_genes on raw expression values, with ctrl_size = 50, n_bins = 25, and random_state = 0. Higher scores indicate stronger expression of an Müller glia-like off-target transcriptional program.

To evaluate bipolar cell type composition shift following retinal regeneration, a Pearson’s chi-square test of independence was performed on raw cell counts across all 27 clusters, accounting for differences in total sample sizes (58,749 control cells vs. 1,389 regenerated cells). Following a significant chi-square result (p<0.0001), post-hoc profiling was conducted by calculating standardized Pearson residuals *r_std_* for each cluster. Shifts in cellular abundance were defined as significant at *r_std_*>1.96 for a=0.05 and highly significant at *r_std_*>3.3 for a=0.001. Positive residuals indicate a relative enrichment in the regenerated retina, while negative residuals denote relative depletion. All calculations were performed in Python using the scipy.stats library (Virtanen et al. 2020).

## Key resources table

## Figure legend

**Supplementary Figure 1. Cell type-specific ablation of retinal neurons following light lesion or NMDA injury.** (A,B) TUNEL staining (A) and quantification (B) of TUNEL+ cells in different retinal layers at 1,2,and 3 days post light lesion. (C,D) Immunostaining (C) and quantification (D) of HuC/D+ neurons in different retinal layers in unlesioned and at 2 and 3 days post light lesion. (One-way ANOVA, GCL p=0.1797; INL: p=0.5348) (E,F) TUNEL staining (E) and quantification (F) of TUNEL+ cells in different retinal layers at 1, 3 hours and 5 days post NMDA injury. ONL: outer nuclear layer; INL: inner nuclear layer; GCL ganglion cell layer. Scale bars 50um.

**Supplementary Figure 2. Validation for specificity and temporal analysis of Tg(mmp9:creERT2; Olactb:loxp-dsRed-loxp-eGFP) lineage tracing line after light lesion.** (A,B) Experimental paradigm and retinal cross sections immunostained for eGFP following 4-hydroxitamoxifen (4-OHT) treatment without retinal injury (A) or light lesion without 4-OHT treatment (B). (C) Immunostaining for Müller glial marker, glutamine synthetase (GS), at 3 days post light lesion. (D,E) Immunostaining for cone (Zpr1, D) and rod (Zpr3, E) photoreceptor markers at 7 days post light lesion. Scale bars 50 um; ONL: outer nuclear layer; INL: inner nuclear layer: GCL: ganglion cell layer.

**Supplementary Figure 3. Immunohistochemical validation of regenerated retinal neurons following NMDA damage.** (A,B) Representative retinal images immunostained for eGFP and rod (Zpr3), cone (Zpr1), bipolar cells (Cabp5), amacrine cells (HuC/D), horizontal cells and retinal ganglion cells (Rbpms) at 14 days post NMDA injury. NMDA: N-methyl-D-aspartate; dpi: days post injection; ONL: outer nuclear layer; INL: inner nuclear layer: GCL: ganglion cell layer. Scale bar 50 um.

**Supplementary Figure 4. Differential gene expression analysis of control and regenerated neurons.** (A-C) Volcano plots representing differentially expressed genes in amacrine cells (A), bipolar cells (B), and horizontal cells (C). (D-F) Boxed plots of selected genes in amacrine cells (D), bipolar cells (E), and horizontal cells (F). LD: light lesion; NMDA: N-methyl-D-aspartate; BC: bipolar cells; AC: amacrine cells; HC: horizontal cells.

**Supplementary Figure 5. ScRNA-Seq subclustering analysis of amacrine cell subtypes across different sample groups (**A) Dot plot showing expression pattern of marker genes across 45 distinct amacrine cell clusters. (B,C) UMAP plot of selected genes unique to a cluster. (D, E) Proportion of cells in 45 amacrine subtypes in unlesioned (D), regenerated (E) retinal samples. (F) Retinal cross section immunostained for GABAergic (Gad65/67), Cholinergic (Chat), and dopaminergic (TH) markers 14 days following NMDA injury. Scale bar 50um. ONL: outer nuclear layer; INL: inner nuclear layer; GCL: ganglion cell layer.

**Supplementary Figure 6. ScRNA-Seq subclustering analysis of bipolar cell subtypes across different sample groups (**A) Dot plot showing expression pattern of marker genes across 27 distinct bipolar cell clusters. (B) UMAP plots of selected genes unique to clusters, 0, 15, and 19. (C) UMAP-plot of immature gene module scores in bipolar cell subtypes. (D) UMAP plots of selected immature marker genes, nfixa and nfixb in bipolar cell subtypes. (E,F) Retinal cross sections immunostained for aPKC (E) and Chat (F) at 14 days post NMDA injection. Scale bars 20 um. ONL: outer nuclear layer; INL: inner nuclear layer: GCL: ganglion cell layer.

**Supplementary Figure 7. Displaced retinal ganglion cells in regenerated retinas. (**A,B) UMAP plots of selected RGC subtype-specific genes. (C,D) Retinal cross section (C) and flat-mount retinas (D) immunostained for Rbpms in control and regenerated animals. (E,F) Higher magnification images of flat-mount retinas immunostained for Rbpms 14 days following light lesion (E) and NMDA injection (F). Scale bars: C,D 200 um; E,F 10 um. D: dorsal; V: ventral; N: nasal; T: temporal; ONL: outer nuclear layer; INL: inner nuclear layer: GCL: ganglion cell layer.

**Supplementary Figure 8. Restoration of precise retinotectal projection.** (A) 3D rendering of contralateral tectum immunostained for eGFP at 14 days post NMDA injection into the right eye, imaged by lightsheet microscopy. (B) Flat-mount ipsilateral tectum immunostained for eGFP at 14 days following NMDA injection into the right eye. (C) Flat-mount tectum immunostained for eGFP at 14 days following light lesion. (D) Cross section of tectum at 14 days following light lesion. Scale bar 200 um. on: optic nerve; a; anterior; p: posterior; SM: Stratum Marginale; SO: Stratum Opticum; SFGC: Stratum Fibrosum Greseum Superficale; SAC: Stratum Griseum Centrale; SPV: Stratum Album Centrale.

Table 1. Differential gene expression analysis of retinal cell clusters (supports Figure 2)

Table 2. Differentially expressed genes between control and regenerated retinal cell types (supports Figure 3)

Table 3. Cluster-specific marker genes across subtypes of amacrine cells (supports Figure 4) Table 4. Cluster-specific marker genes across subtypes of bipolar cells (supports Figure 5)

Table 5. Cluster-specific marker genes across subtypes of retinal ganglion cells (supports Figure 6) Table 6. Key reagents used in the study

