## Supplementary Figures for "Müller glia–mediated regeneration restores neuronal diversity and circuit organization in the adult zebrafish retina"

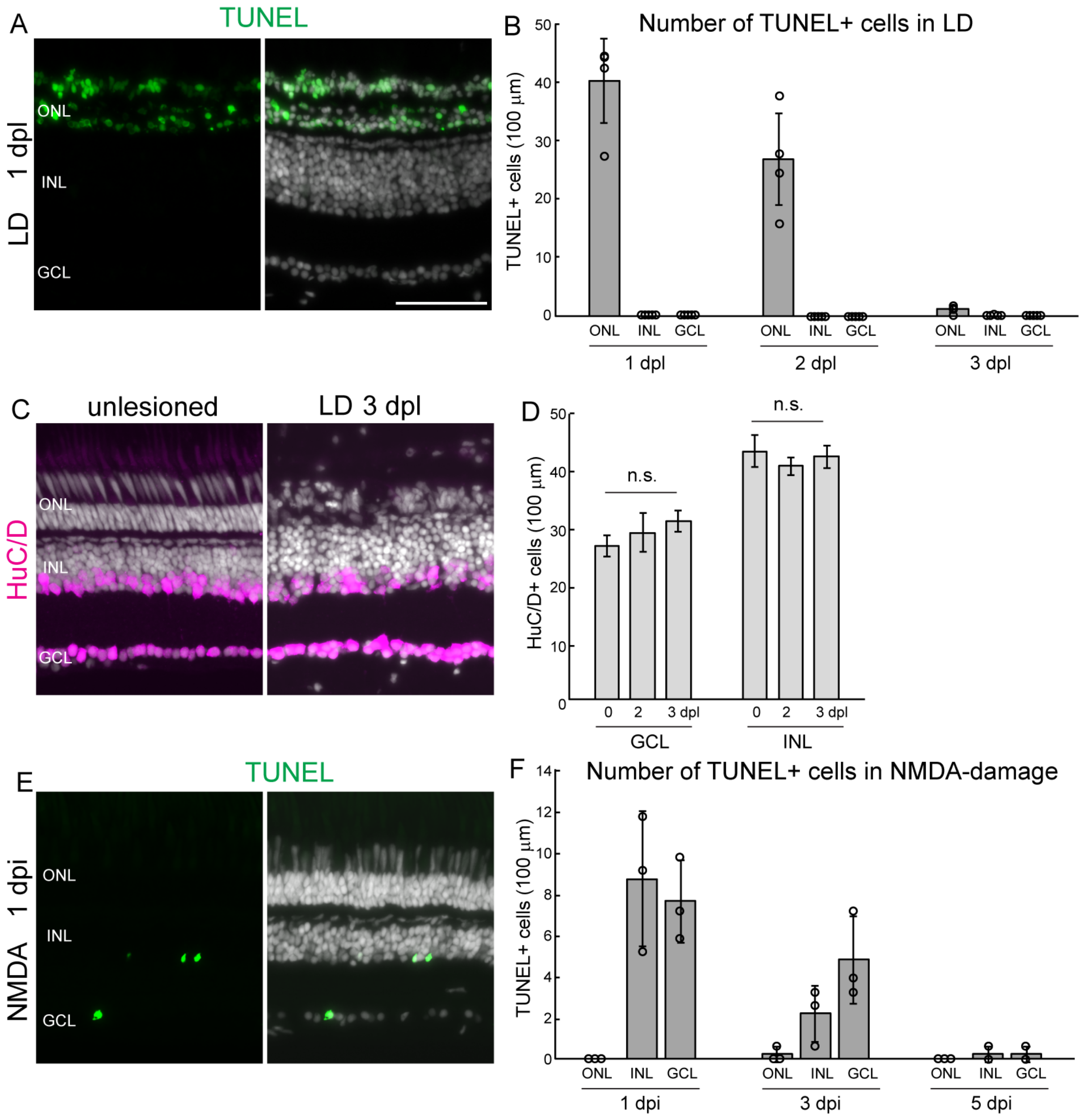


**Supplementary Figure 1. Cell type-specific ablation of retinal neurons following light lesion or NMDA injection.** (A,B) TUNEL staining (A) and quantification (B) of TUNEL+ cells in different retinal layers at 1,2,and 3 days post light lesion. (C,D) Immunocytochemistry (C) and quantification (D) of HuC/D+ neurons in different retinal layers in unlesioned and at 2 and 3 days post light lesion. (One-way ANOVA, GCL p=0.1797; INL: p=0.5348) (E,F) TUNEL staining (E) and quantification (F) of TUNEL+ cells in different retinal layers at 1, 3 hours and 5 days post NMDA injection. ONL: outer nuclear layer; INL: inner nuclear layer; GCL ganglion cell layer. Scale bars 50um.

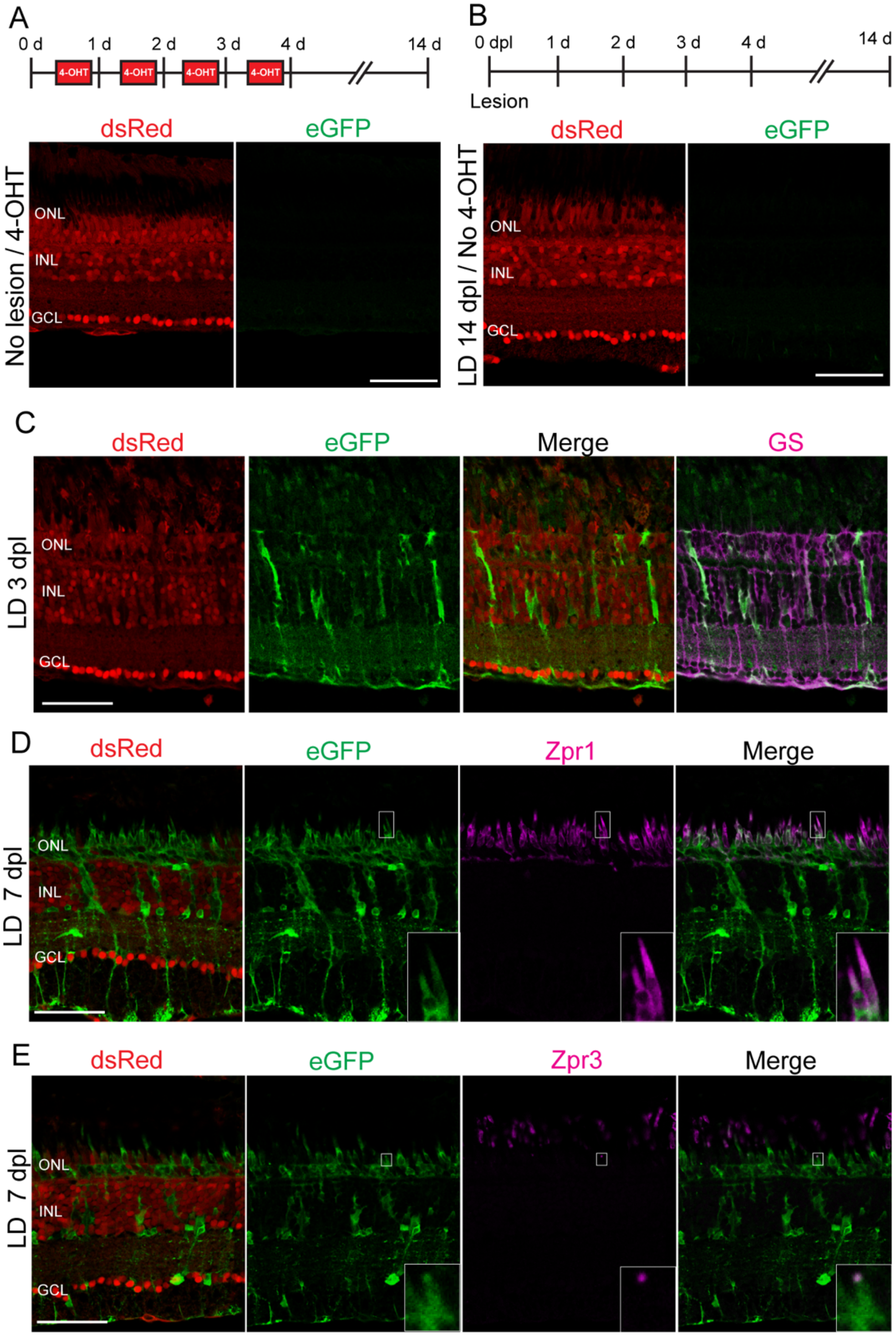


**Supplementary Figure 2. Validation for specificity and temporal analysis of Tg(mmp9:creERT2; Olactb:loxp-dsRed-loxp-eGFP) lineage tracing line after light lesion.** (A,B) Experimental paradigm and retinal cross sections immunostained for eGFP following 4-hydroxitamoxifen (4-OHT) treatment without retinal injury (A) or light lesion without 4-OHT treatment (B). (C) Immunocytochemistry for Müller glial marker, glutamine synthetase (GS), at 3 days post light lesion. (D,E) Immunocytochemistry for cone (Zpr1, D) and rod (Zpr3, E) photoreceptor markers at 7 days post light lesion. Scale bars 50 um; ONL: outer nuclear layer; INL: inner nuclear layer: GCL: ganglion cell layer.

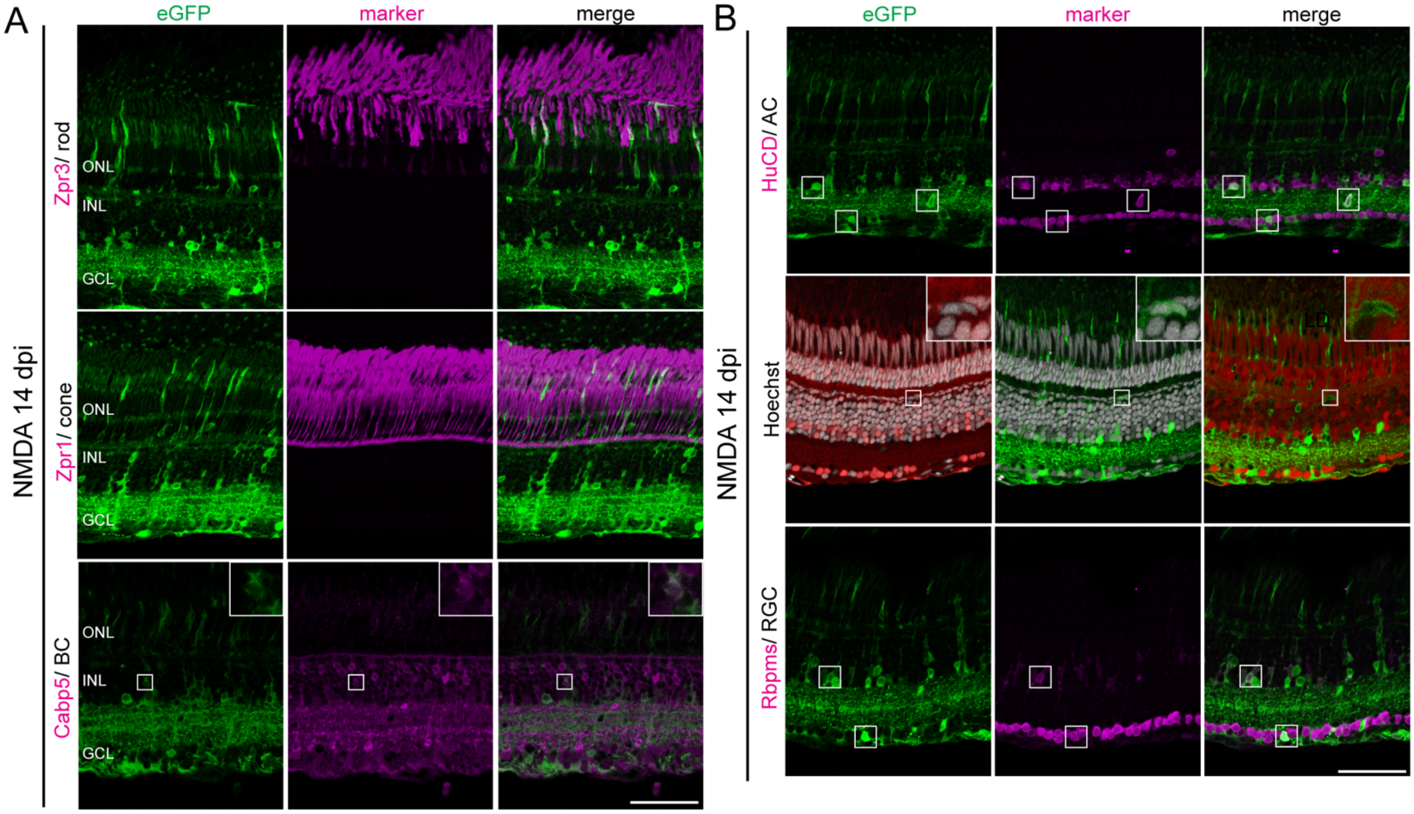


**Supplementary Figure 3. Immunohistochemical validation of regenerated retinal neurons following NMDA damage.** (A,B) Representative retinal images immunostained for eGFP and rod (Zpr3), cone (Zpr1), bipolar cells (Cabp5), amacrine cells (HuC/D), horizontal cells and retinal ganglion cells (Rbpms) at 14 days post NMDA injection. NMDA: N-methyl-D-aspartate; dpi: days post injection; ONL: outer nuclear layer; INL: inner nuclear layer: GCL: ganglion cell layer. Scale bar 50 um.

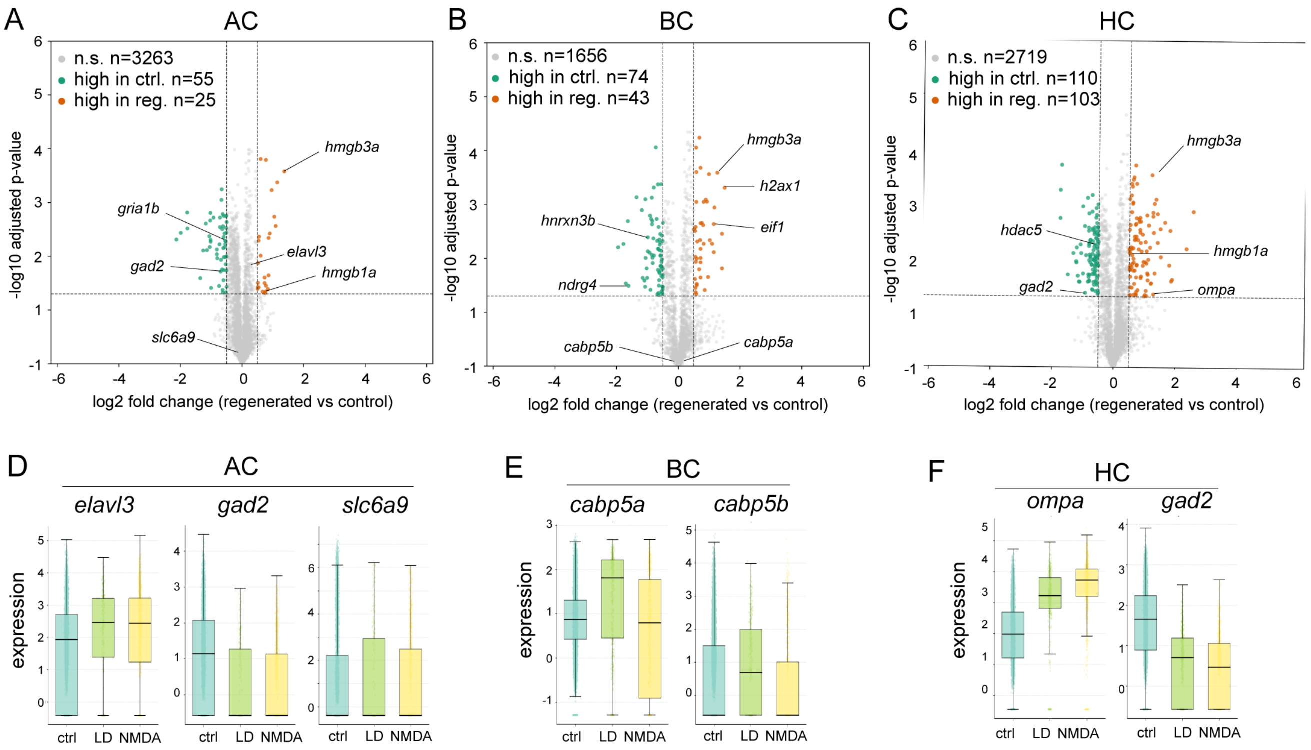


**Supplementary Figure 4. Differential gene expression analysis of control and regenerated neurons.** (A-C) Volcano plots representing differentially expressed genes in amacrine cells (A), bipolar cells (B), and horizontal cells (C). (D-F) Boxed plots of selected genes in amacrine cells (D), bipolar cells (E), and horizontal cells (F). LD: light lesion; NMDA: N-methyl-D-aspartate; BC: bipolar cells; AC: amacrine cells; HC: horizontal cells.

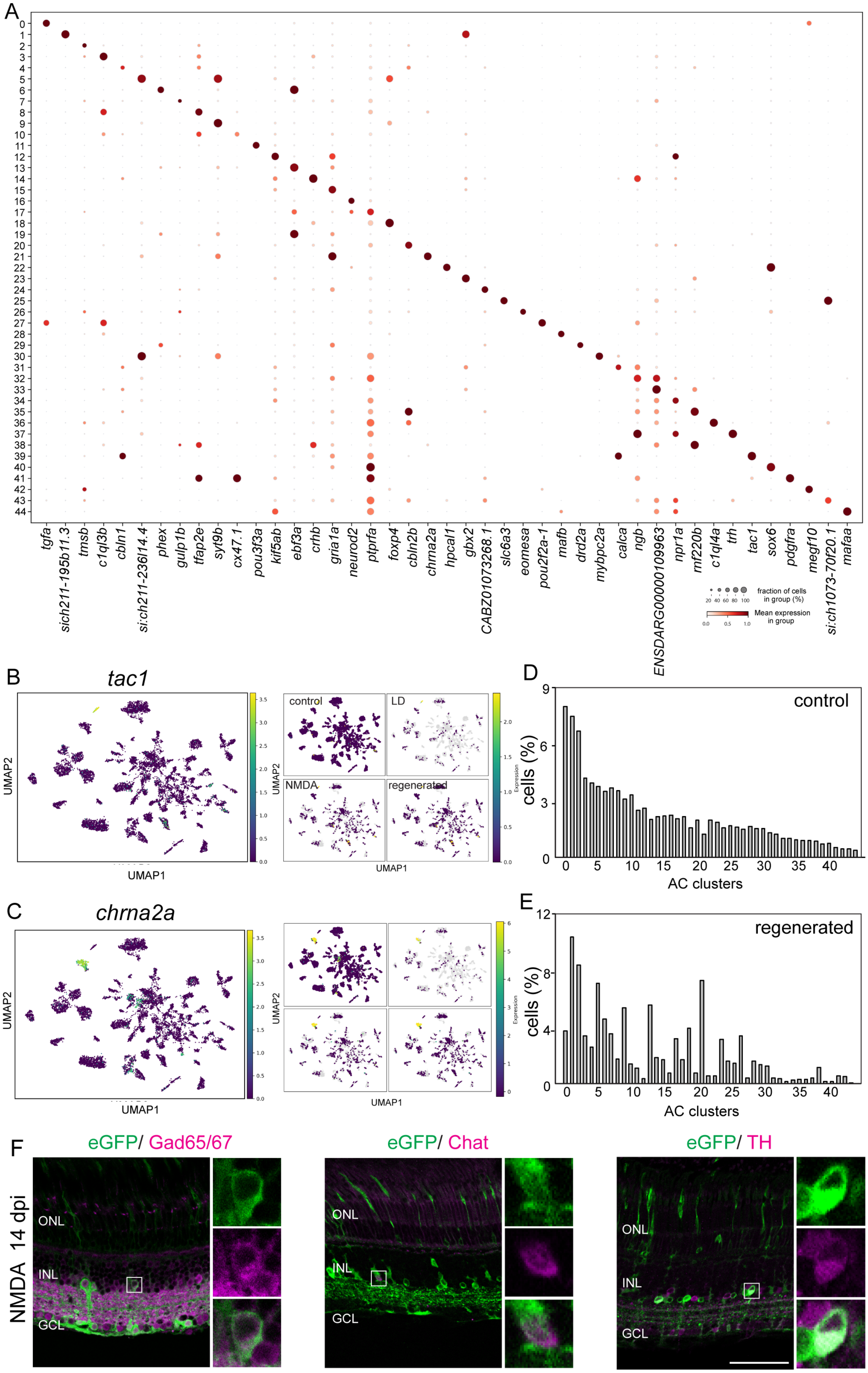

A
tgfa
sich211-195b11.3
tmsb
c1ql3b
cbln1
si:ch211-236l14.4
phex
gulp1b
tfap2e
syt9b
cx47.1
pou3f3a
kif5ab
ebf3a
crhb
gria1a
neurod2
ptprfa
foxp4
cbln2b
chrna2a
hpcal1
gbx2
CABZ01073268.1
slc6a3
eomesa
pou2f2a-1
mafb
drd2a
mybpc2a
calca
ngb
ENSDARG00000109963
npr1a
mf220b
c1ql4a
trh
tac1
sox6
pdgfra
megf10
si:ch1073-70f20.1
mafaa
fraction of cells in group (%)
Mean expression in group
B
tac1
UMAP2
UMAP1
control
LD
NMDA
regenerated
Expression
C
chrna2a
D
control
cells (%)
AC clusters
E
regenerated
F
eGFP/ Gad65/67
eGFP/ Chat
eGFP/ TH
NMDA 14 dpi
ONL
INL
GCL

**Supplementary Figure 5. ScRNA-Seq subclustering analysis of amacrine cell subtypes across different sample groups**. **(**A) Dot plot showing expression pattern of marker genes across 45 distinct amacrine cell clusters. (B,C) UMAP plot of selected genes unique to a cluster. (D, E) Proportion of cells in 45 amacrine subtypes in unlesioned (D), regenerated (E) retinal samples. (F) Retinal cross section immunostained for GABAergic (Gad65/67), Cholinergic (Chat), and dopaminergic (TH) markers 14 days following NMDA injection. Scale bar 50um. ONL: outer nuclear layer; INL: inner nuclear layer; GCL: ganglion cell layer.

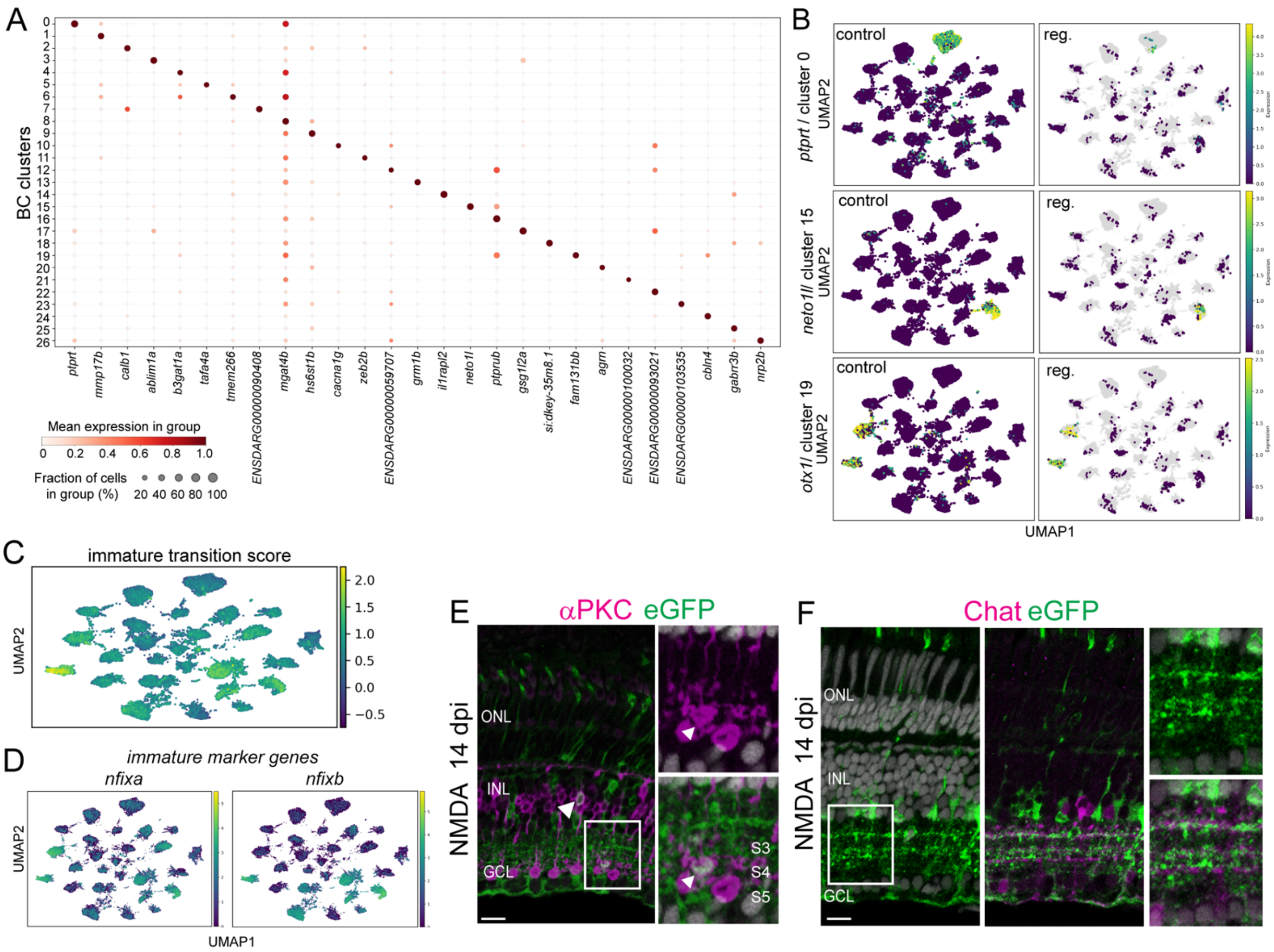


**Supplementary Figure 6. ScRNA-Seq subclustering analysis of bipolar cell subtypes across different sample groups**. (A) Dot plot showing expression pattern of marker genes across 27 distinct bipolar cell clusters. (B) UMAP plots of selected genes unique to clusters, 0, 15, and 19. (C) UMAP-plot of immature gene module scores in bipolar cell subtypes. (D) UMAP plots of selected immature marker genes, nfixa and nfixb in bipolar cell subtypes. (E,F) Retinal cross sections immunostained for aPKC (E) and Chat (F) at 14 days post NMDA injection. Scale bars 20 um. ONL: outer nuclear layer; INL: inner nuclear layer: GCL: ganglion cell layer.

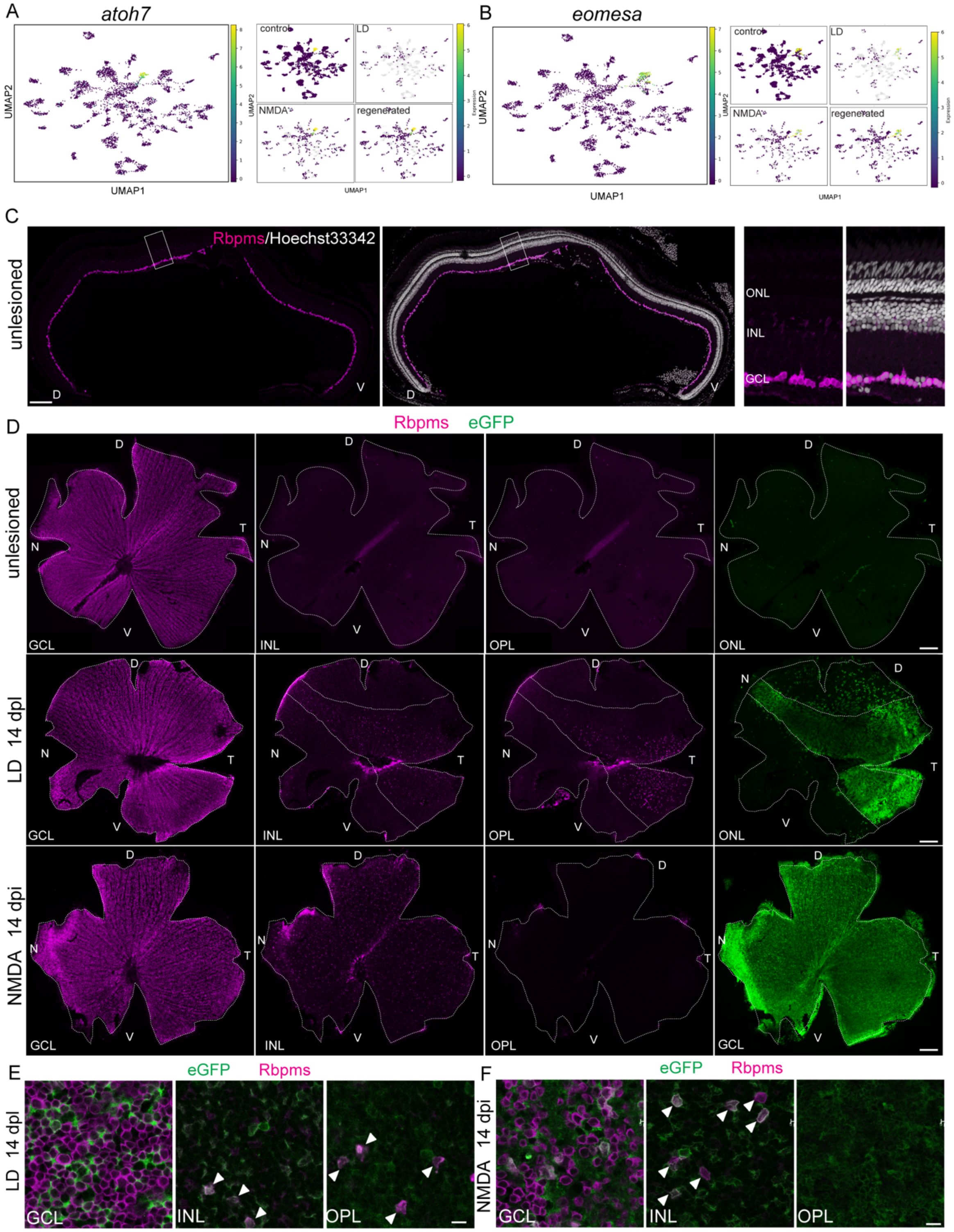


**Supplementary Figure 7. Displaced retinal ganglion cells in regenerated retinas. (**A,B) UMAP plots of selected RGC subtype-specific genes. (C,D) Retinal cross section (C) and flat-mount retinas (D) immunostained for Rbpms in control and regenerated animals. (E,F) Higher magnification images of flat-mount retinas immunostained for Rbpms 14 days following light lesion (E) and NMDA injection (F). Scale bars: C,D 200 um; E,F 10 um. D: dorsal; V: ventral; N: nasal; T: temporal; ONL: outer nuclear layer; INL: inner nuclear layer: GCL: ganglion cell layer.

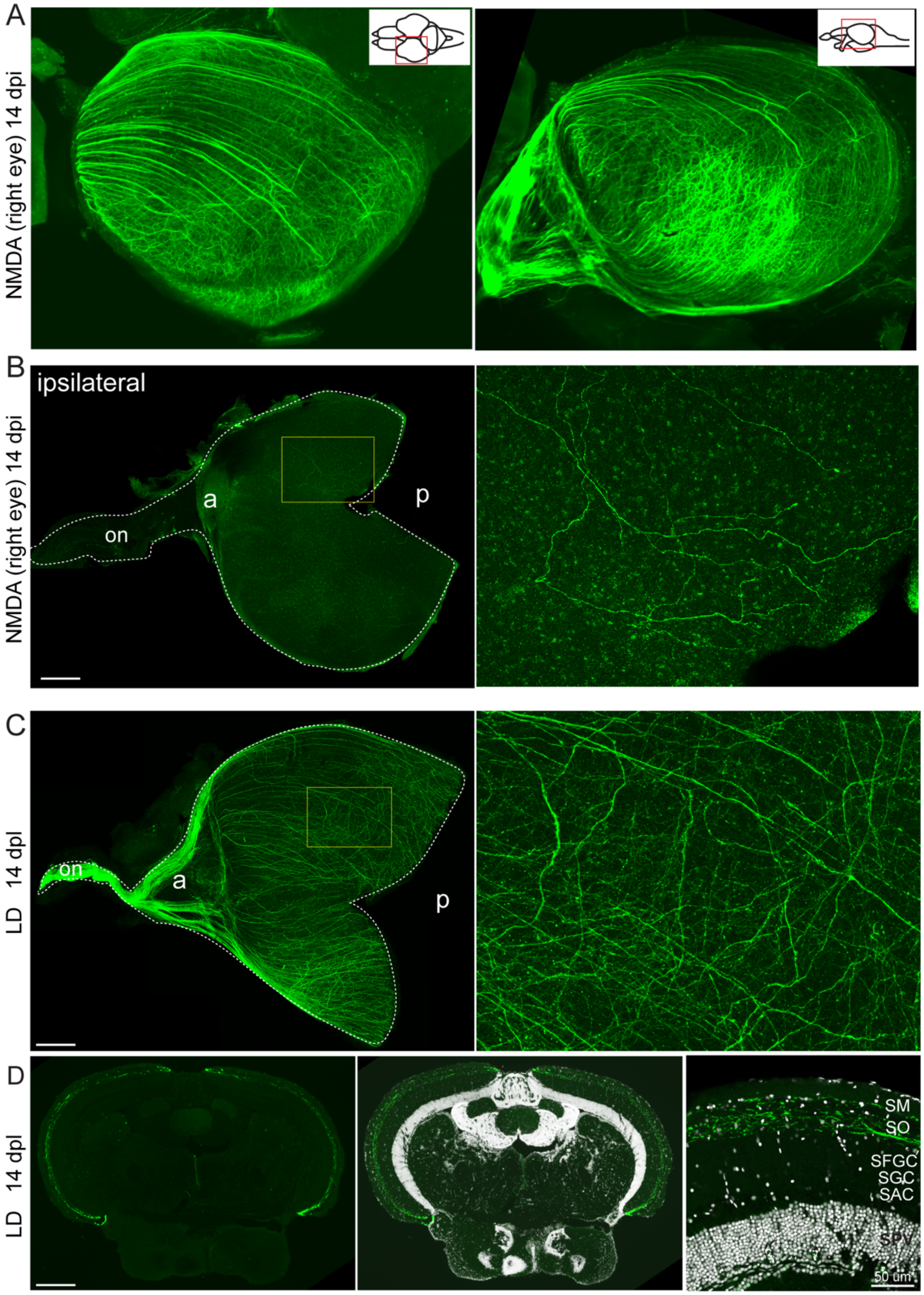


**Supplementary Figure 8. Restoration of precise retinotectal projection.** (A) 3D rendering of contralateral tectum immunostained for eGFP at 14 days post NMDA injection into the right eye, imaged by lightsheet microscopy. (B) Flat-mount ipsilateral tectum immunostained for eGFP at 14 days following NMDA injection into the right eye. (C) Flat-mount tectum immunostained for eGFP at 14 days following light lesion. (D) Cross section of tectum at 14 days following light lesion. Scale bar 200 um. on: optic nerve; a; anterior; p: posterior; SM: Stratum Marginale; SO: Stratum Opticum; SFGC: Stratum Fibrosum Greseum Superficale; SAC: Stratum Griseum Centrale; SPV: Stratum Album Centrale.
